# Toward an optimal contraception dosing strategy

**DOI:** 10.1101/2022.04.04.486926

**Authors:** Brenda Lyn A. Gavina, Aurelio A. de los Reyes, Mette S. Olufsen, Suzanne Lenhart, Johnny T. Ottesen

## Abstract

Anovulation refers to a menstrual cycle characterized by the absence of ovulation. Exogenous hormones such as synthetic progesterone and estrogen have been used to attain this state to achieve contraception. However, large doses are associated with adverse effects such as increased risk for thrombosis and myocardial infarction. This study utilizes optimal control theory on a modified menstrual cycle model to determine the minimum total exogenous estrogen/progesterone dose, and timing of administration to induce anovulation. The mathematical model correctly predicts the mean daily levels of pituitary hormones *LH* and *FSH*, and ovarian hormones *E*_2_, *P*_4_, and *Inh* throughout a normal menstrual cycle and reflects the reduction in these hormone levels caused by exogenous estrogen and/or progesterone. Results show that it is possible to reduce the total dose by 92% in estrogen monotherapy, 43% in progesterone monotherapy, and that it is most effective to deliver the estrogen contraceptive in the mid follicular phase. Finally, we show that by combining estrogen and progesterone the dose can be lowered even more. These results may give clinicians insights into optimal formulations and schedule of therapy that can suppress ovulation.

**AUTHOR SUMMARY:** Hormonal contraceptives composed of exogenous estrogen and/or progesterone are commonly administered artificial means of birth control. Despite many benefits, adverse side effects associated with high doses such as thrombosis and myocardial infarction, cause hesitation to usage. Our study presents an improved mathematical model for hormonal control of the menstrual cycle and applies optimal control theory to minimize total exogenous estrogen and/or progesterone dose, and determine timing of administration that lead to contraception. We observe a reduction in dosage of about 92% in estrogen monotherapy and 43% in progesterone monotherapy. Our simulations show that it is most effective to deliver the estrogen contraceptive in the mid follicular phase. In addition, we illustrate that combination therapy significantly lower doses further. Our findings may give clinicians insights into optimal dosing scheme for contraception.

## INTRODUCTION

The female’s reproductive life spanning approximately 39 years from age of 12.5 until 51 is governed by the menstrual cycle [1], a cyclic process regulated by the endocrine system. A normal menstrual cycle involves ovarian follicular development, ovulation, and luteinization influenced by the hormones gonadotropin-releasing hormone (GnRH), luteinizing hormone (*LH*), follicle-stimulating hormone (*FSH*), estradiol (*E*_2_), progesterone (*P*_4_), and inhibin (*Inh*), which are produced in the hypothalamus, pituitary, and ovaries [2, 3]. During this cycle, the pituitary and ovarian hormones fluctuate. Unusual concentrations of these hormones lead to abnormal cycles. For instance, low levels of *LH, FSH*, and *E*_2_ cause anovulation [3]. Anovulation is an abnormal menstrual cycle characterized by the absence of the ovulation process [4].

Numerous previous modeling studies have examined the menstrual cycle, how it is formed and how it can be altered. The most significant body of work stems from Selgrade et al. [5–13]. These studies start by developing a continuous menstrual cycle model and fitting it to data. This model consists of two parts tracking the pituitary [6] and ovarian [5] portions of the cycle. The full model developed by Harris et al. [7, 8, 12] is formulated as an autonomous nonlinear system of 13 delay differential equations (DDEs) merging the pituitary and ovarian components. The model, consisting of positive and negative feedback relationships of the pituitary and ovarian hormones, was fitted to data from McLachlan et al. [14] reporting daily blood concentrations of *LH, FSH, E*_2_, *P*_4_, and *Inh* averaged from data for 33 normally cycling women. Pasteur [9] expanded this model by distinguishing the effect of inhibin A (*Inh A*) and inhibin B (*Inh B*). The inclusion of the two forms of inhibin provides a more realistic representation of the human menstrual cycle since these hormones are active in different phases of the cycle [9]. The model was further developed in the study by Margolskee et al. [11] who reduced the number of delays to one (delay in inhibin) and fitted it to data by Welt et al. [15], which consist of daily mean blood levels of *LH, FSH, E*_2_, *P*_4_, and *Inh A* from 23 normally cycling women. Most recently, Wright et al. [13] added autocrine mechanisms to the Margolskee model to describe the influence of exogenous estrogen and/or progesterone in perturbing a normal cycle to contraceptive state.

In addition to work by Selgrade and collaborators, Chen et al. [16] introduced a simple model with three delay differential equations describing the hormonal interactions of the human menstrual cycle along the hypothalamus–pituitary–ovaries axis. Reinecke et al. [17, 18] studied the pulsatile release of GnRH in the hypothalamus using a stochastic process, and Röblitz et al. [19] utilized ordinary differential equations (ODEs) to examine the interactions between GnRH, pituitary, and ovarian hormones which shortened computational time.

These menstrual cycle models and their modifications, all fitted to biological data consisting of blood levels of pituitary and ovarian hormones from normally cycling women, have been used to study fertility [7, 9, 12, 19], reproductive diseases [16], and contraception [13, 17–19]. For instance, Pasteur’s model showed that the dosage of exogenous estrogen varies inversely as the amplitudes of hormone level fluctuations, while Chen’s model simulated treatment of uterine myomas with GnRH analogues.

Contraception is achieved through natural and/or artificial means. Artificial methods include hormonal, barrier, permanent, and long-acting reversible contraceptives. Today, contraception is most commonly achieved by taking a daily pill, though this method is rapidly being replaced by injectables and implants [20]. Independent of the administration method, almost all hormonal contraceptives including exogenous progesterone and/or estrogen act by blocking ovulation, changing cervical mucus, which hinders sperm transport and/or modifying endometrium which prevents implantation [21]. Aside from contraceptive benefit, suppression of ovulation can alleviate negative premenstrual symptoms [22–24]. For example, Hammarbäck et al. [23] showed that cyclical negative premenstrual symptoms such as irritability and breast tenderness disappear in anovulatory cycles. In addition, Yonkers et al. [24] found that administration of GnRH analogues, exogenous estrogen, and certain oral contraceptives at doses which inhibit ovulation is effective in reducing the symptoms. Most literature studies [13, 18, 19] focus on the administration of exogenous hormones such as estrogen, progesterone, and GnRH analogues to inhibit ovulation. For example, Reinecke’s detailed DDE model included a numerical study on the parts of body affected by estrogen and/or progesterone hormonal contraceptives. This model compared results from administering the contraceptive continuously at a constant rate and at certain time points. Roblitz ODE model compares the effect of administering a single and multiple dose of Nafarelin (a GnRH agonist) and Cetrorelix (a GnRH antagonist known to impede ovulation in in-vitro fertilization treatment), and Wright’s model simulated exogenous estrogen and/or progesterone doses concluding that by combining the two effectors the dose can be lowered significantly. These studies all explored the effect of exogenous hormones for inducing anovulation but did not examine if optimal dosing can reduce the dose further. With rapid advance in implants and injections providing continuous administration there is great potential to implement new patient-specific minimizing dosing schemes. To our knowledge, our work is the first to use modeling to study timing of dosing thereby minimizing the dose even more. As implants become more common, results from this study have potential to provide contraception to more women, in particular since lower doses also decrease the risks for adverse side effects such as venous thromboembolism and myocardial infarction associated with high doses of hormonal contraceptives [3, 25–27].

While optimal control theory has not been used for contraception, the theory has long tradition in biology to find strategies that optimize an outcome. In [28], the theory is applied to a system of ODEs to determine a scheme for delivery of insulin and glucagon that regulates blood glucose level in diabetes patients. The study [29] utilized optimal control to develop treatment protocol for tumor stabilization for prostate cancer. The current study employs optimal control theory to a modification of the model in Margolskee et al. [11] to determine the minimum total dosage of exogenous estrogen/progesterone, and timing of administration that induce anovulation. The mathematical model includes mechanisms depicting the contraceptive effect of exogenous progesterone on the menstrual cycle. This new model for contraception calibrated to the data extracted from Welt et al. [15] predicts the daily levels of pituitary hormones *LH* and *FSH*, and ovarian hormones *E*_2_, *P*_4_, and *Inh* averaged during a normal menstrual cycle. The model output also predicts reduction in pituitary and ovarian hormone levels caused by exogenous estrogen and/or progesterone observed by Obruca et al. [30] and Deb et al. [31]. Simulation results show that the dosage may be reduced by 92% in estrogen monotherapy, 43% in progesterone monotherapy, to suppress ovulation. Simulations also agree with biological literature that in monotherapy, administration of estrogen in the mid follicular phase is effective at preventing ovulation. Lastly, numerical experiments show that by combining estrogen and progesterone the dose can be reduced even further. The results of this study may aid in identifying the minimum dose and treatment schedule that cause anovulation.

## METHODS

This section describes the normal and anovulatory menstrual cycle, data, the mathematical model, parameter estimation, and optimal control method. Tables I and V list the state variables and parameters in the mathematical model.

**TABLE I.** Menstrual cycle state variables and initial conditions.

| State variable | Description | Initial condition | Unit | Reference |
| --- | --- | --- | --- | --- |
| $RP_{LH}$ | Amount of reserve pool $LH$ | 167.57 | IU | estimated |
| $LH$ | $LH$ blood concentration | 11.81 | IU/L | estimated |
| $RP_{FSH}$ | Amount of reserve pool $FSH$ | 14.48 | IU | estimated |
| $FSH$ | $FSH$ blood concentration | 11.41 | IU/L | estimated |
|  | Amount of active follicular mass |  |  |  |
| $RcF$ | in the recruited follicular stage | 2.10 | ng | estimated |
| $GrF$ | in the growing follicular stage | 4.12 | ng | estimated |
| $DomF$ | in the dominant follicular stage | 0.46 | ng | estimated |
| $Sc_1$ | during ovulation | 1.06 | ng | estimated |
| $Sc_2$ | during luteinization | 1.67 | ng | estimated |
|  | Amount of active luteal mass |  |  |  |
| $Lut_1$ | in the first luteal stage | 4.16 | ng | estimated |
| $Lut_2$ | in the second luteal stage | 13.03 | ng | estimated |
| $Lut_3$ | in the third luteal stage | 16.48 | ng | estimated |
| $Lut_4$ | in the fourth luteal stage | 10.29 | ng | estimated |

### The normal menstrual cycle

The normal menstrual cycle (shown in Figure 1) for an adult female has an average length of 28 days. It has two stages, the follicular phase and the luteal phase. Through hormones, the hypothalamus, pituitary, and ovaries interact to regulate the menstrual cycle [2, 3].

**FIG. 1.**
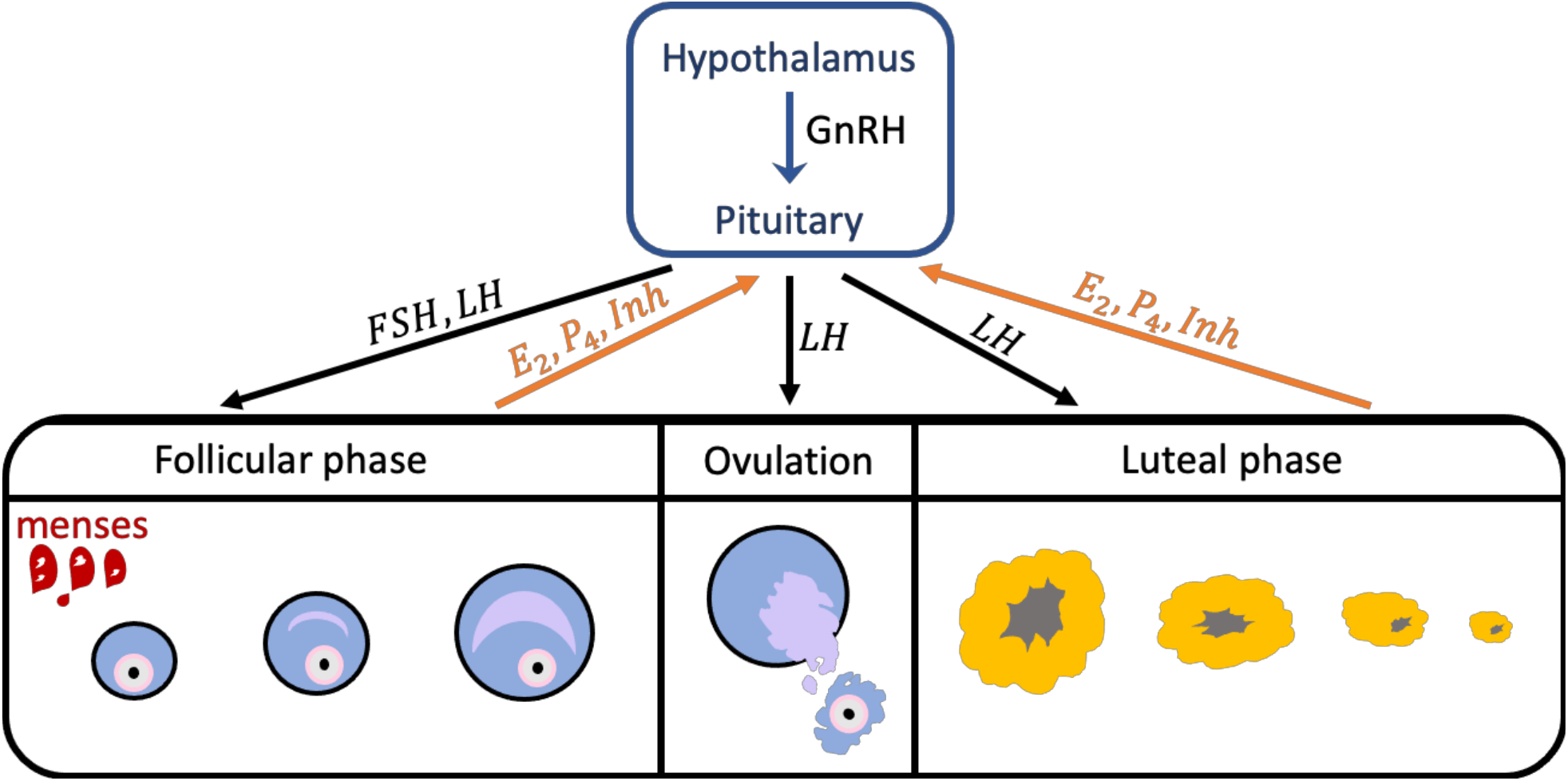
The Follicular and Luteal Phases of the menstrual cycle. The figure shows the transition of a follicle, from its growth in the follicuar phase to its rupture during ovulation, as well as its transformation to a corpus luteum and degradation in the luteal phase. The blue arrow indicates the control of the hypothalamus in the secretion of pituitary hormones. The black arrows represent the influence of the pituitary system on the ovarian system through *FSH* and *LH*, and the orange arrows show the response of the ovarian system through *E*_2_, *P*_4_, and *Inh*.

During the menstrual cycle, the hypothalamus produces pulses of GnRH which control the pituitary’s secretion of the gonadotropins *FSH* and *LH*. At the beginning of the follicular phase (the start of menstruation or menses), *FSH* rises and causes the recruitment of a group of immature follicles. As these follicles develop through the stimulation of the gonadotropins *FSH* and *LH*, they increase secretion of *E*_2_ (see Figure 2). Follicular development is indicated by an enlargement of oocyte (immature egg), multiplication, and transformation of granulosa cells (structure that surrounds the oocyte) to a cuboidal shape, and formation of small gap junctions, which enable nutritional, metabolite, and signal interchange between the granulosa cells and oocyte [3]. Toward the end of the follicular stage, one dominant follicle continues to growth while the rest of the follicles become atretic. As the dominant follicle grows, the *E*_2_ level increases to a maximum value prompting an *LH* surge. The *LH* surge stimulates ovulation, releasing the egg from the dominant follicle. The ruptured dominant follicle then transforms to a corpus luteum. As the corpus luteum grow, *P*_4_ and *Inh* production is increased. *P*_4_ and *Inh* inhibit the synthesis of *LH* and *FSH*. If fertilization does not occur, the corpus luteum degrades removing the inhibition on *LH* and *FSH*. Consequently, levels of *FSH* and *LH* rise and the menstrual cycle repeats [2, 3].

**FIG. 2.**
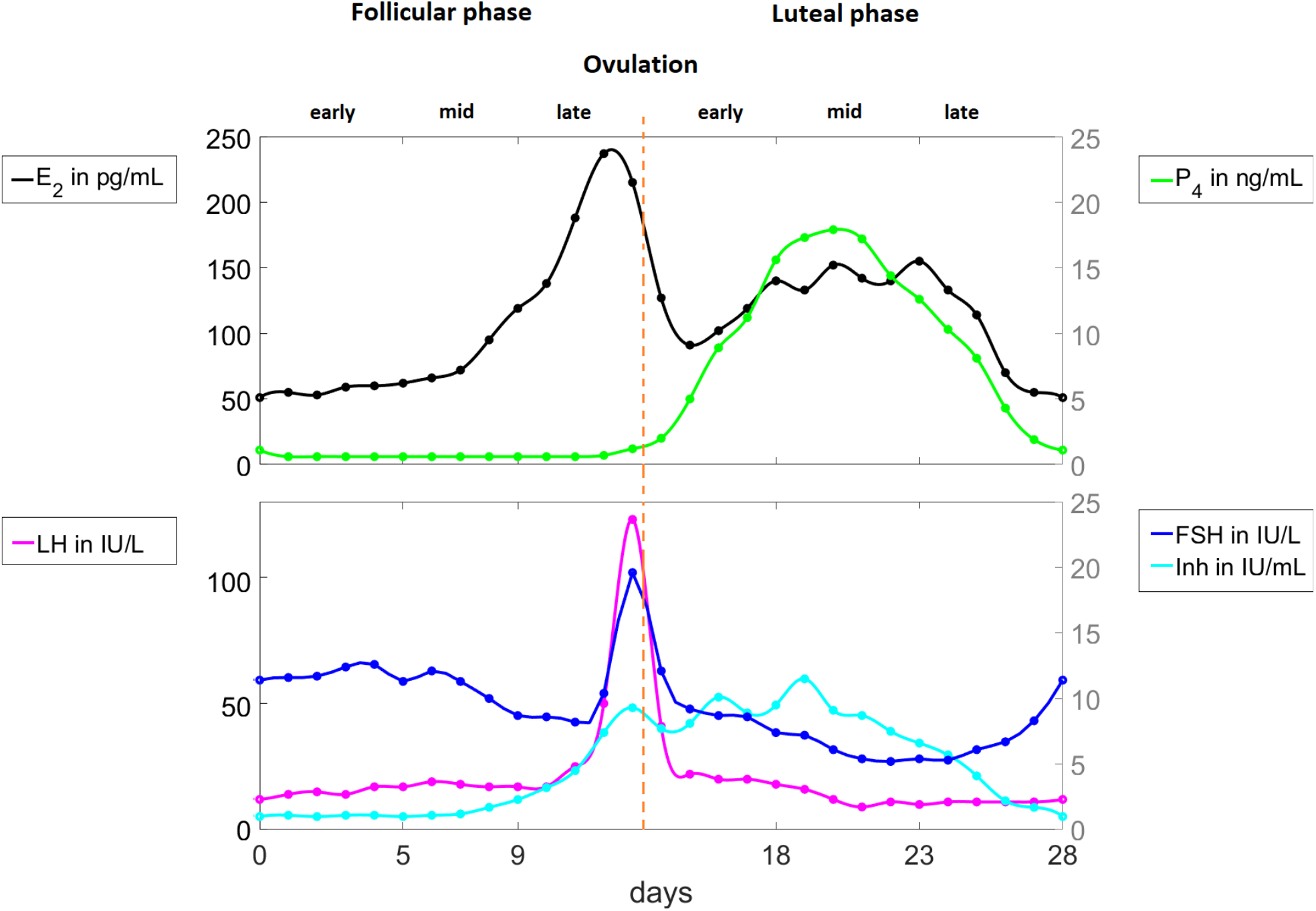
Pituitary and ovarian hormone levels in a normal menstrual cycle. Data digitized from the study by Welt et al. [15] is interpolated by cubic splines. The hormones *LH, FSH*, and *E*_2_ peak in the late follicular phase while *P*_4_ and Inh reach maximum value in the luteal phase.

### The anovulatory cycle and contraception

Anovulation occurs when the follicle fails to release the egg. This state can be detected by measuring serum progesterone. During a normal menstrual cycle, progesterone is largely produced by the corpus luteum after ovulation. Its levels stay below 2 ng/mL during the follicular phase and peak 7 to 8 days after ovulation [3]. Accordingly, a low concentration of this hormone in the luteal phase indicates anovulation or defective luteinization [32]. The menstrual cycle is classified as anovulatory if progesterone concentration remains below 5 ng/mL without an *LH* peak [33]. In fact, disruption at any level of the hypothalamic-pituitary-gonadal axis can result in anovulation. This includes suppression of GnRH, the presence of a pituitary tumor leading to gonadotropin secretory dynamics that fail to stimulate follicular growth, or abnormal estrogen feedback signaling causing inhibition of *FSH* secretion or a low estrogen level which prevent the *LH* surge [3].

Hormonal contraceptives, composed of progesterone alone, or a combination of estrogen and progesterone are widely used artificial means of contraception. They are delivered orally, transdermally, vaginally, via implants, or injections. Estrogen or progesterone alone can cause contraception via anovulation but the combined administration of both hormones significantly enhances effectiveness [34].

In oral steroid contraceptives composed of both exogenous estrogen and progesterone, progesterone induces anovulation by preventing the midcycle rise in *LH* secretion [34, 35]. This is due to progesterone’s suppression of follicular development and gonadotropin secretion [36]. The insufficient *FSH* prevents follicle growth. Consequently, the lack of follicle growth results in an inadequate amount of estradiol inhibiting the *LH* surge [3]. The estrogen component suppresses *FSH* secretion blocking folliculogenesis, stabilizing the endometrium, which minimize bleeding [35, 37].

### Data

The output of the mathematical model without the administration of exogenous hormone is compared to the 28-day data (solid circles in Figure 2) from Welt et al. [15]. These data, extracted from Figure 1 in [15] using the software DigitizeIt version 2.5 [38], comprise mean levels of *E*_2_, *P*_4_, *Inh A, LH*, and *FSH* taken from 23 normally cycling younger women aged 20 to 34 years. Our study employs only *Inh A* since *Inh B* is more significant in studies about reproductive aging.

### Mathematical model of the normal menstrual cycle

To study anovulation, we use the normal menstrual cycle model by Margolskee et al. [11] and induce anovulation via added estrogen and progesterone. The core model, shown in Figure 3, includes the pituitary and ovarian phases. It assumes that

**FIG. 3.**
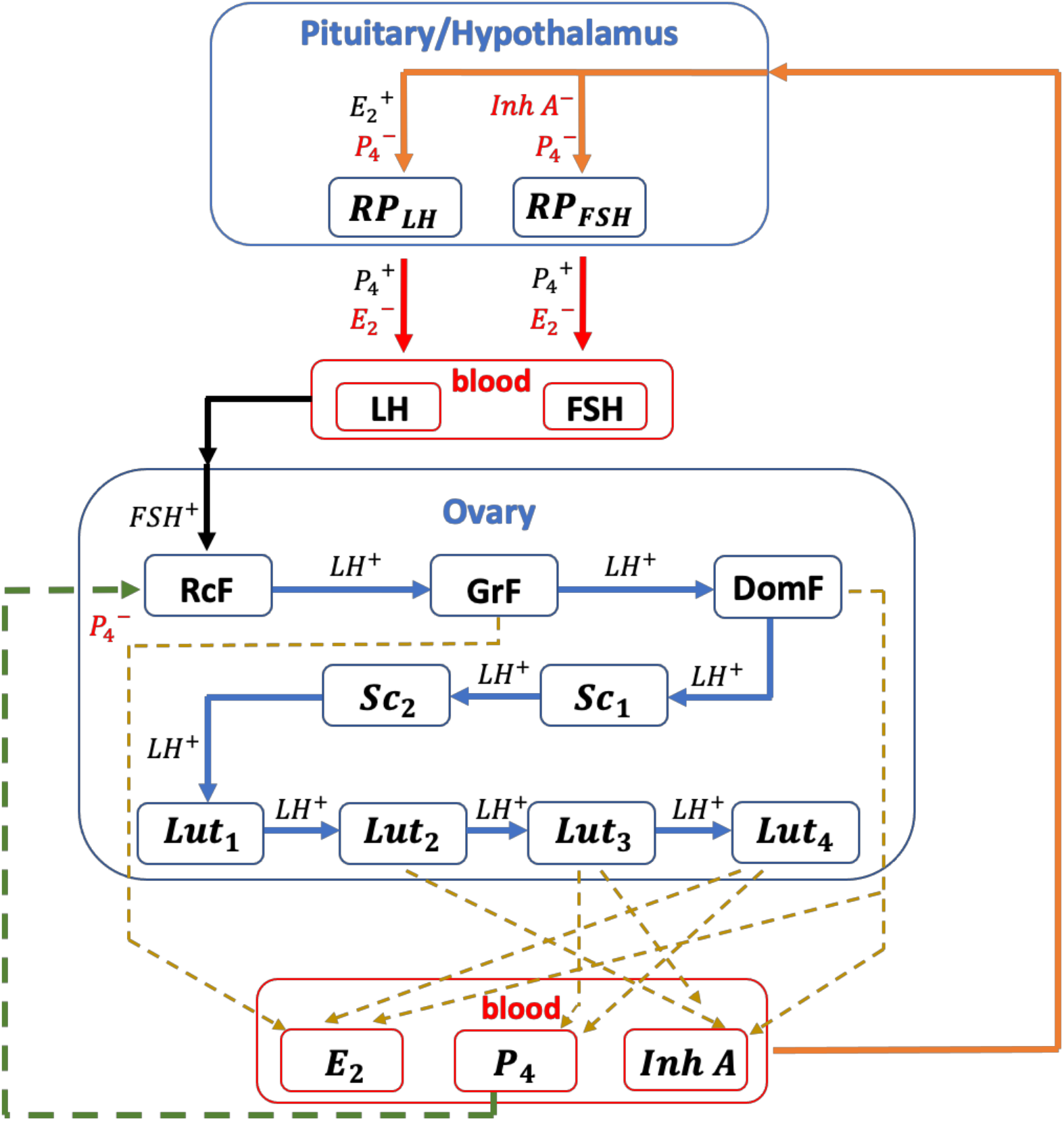
Model diagram. The diagram (adapted from [13]) shows the 13 states (*RP*_*LH*_, *RP*_*F SH*_, *LH, FSH, RcF, GrF, DomF, Sc*_1_, *Sc*_2_, *Lut*_1_, *Lut*_2_, *Lut*_3_, *Lut*_4_) in the menstrual cycle model. *H*^+^ denotes stimulation by hormone *H* while *H*^−^ denotes inhibition. The red arrow presents the release of the pituitary hormone from the reserve pool to the blood. The black arrow indicates the influence of the pituitary system on the ovarian system while the orange arrow denotes the ovary’s feedback. The green dashed arrow denote inhibition of *P*_4_ in the *RcF* stage. The blue arrow desribes the transition of a follicle from one ovarian stage to the next. The gold dashed arrow denotes the state contributing to the ovarian hormone production.

a.1 *LH* and *FSH* synthesis occur in the pituitary,
a.2 *LH* and *FSH* are held in a reserve pool awaiting release into the bloodstream, and
a.3 the follicular/luteal mass undergoes nine ovarian stages of development.

In the model, *RP*_*LH*_(*t*) and *RP*_*FSH*_ (*t*) denote the amount of *LH* and *FSH* in the reserve pool at time *t* days, *LH*(*t*) and *FSH*(*t*) are the blood concentrations of *LH* and *FSH*; *E*_2_(*t*), *P*_4_(*t*), and *Inh*(*t*) denote the blood levels of *E*_2_, *P*_4_, and *Inh*; and *RcF* (*t*), *GrF* (*t*), *DomF* (*t*), *Sc*_1_(*t*), *Sc*_2_(*t*), *Lut*_1_(*t*), *Lut*_2_(*t*), *Lut*_3_(*t*), and *Lut*_4_(*t*) are the masses of active follicular/luteal tissues in the nine ovarian stages: recruited, growing, and dominant follicular stages, ovulation, luteinization, first, second, third, and fourth stages of luteal development.

**The pituitary model** includes the hypothalamus and the pituitary. It predicts the synthesis, release, and clearance of *LH* and *FSH*, and the pituitary’s response to *E*_2_, *P*_4_, and *Inh* (see Figure 3). The direct effect of the ovarian hormones on the pituitary and indirect influence via the hypothalamus are grouped partly because of the complexity of tracking GnRH [6].

The rate of change of *RP*_*LH*_(*t*) in Equation (1) is the difference between two terms. The first term describes the synthesis in the pituitary and the second term represents the release of *LH* into the blood. A Hill function is used in the synthesis term of *LH* to predict the strong stimulatory effect for *E*_2_ levels above the threshold level *Km*_*LH*_. *P*_4_ inhibits the production but bolsters the *LH* release into the bloodstream. *E*_2_ inhibits the release of *LH*. The change in *LH*(*t*) in Equation (2) is affected by the *LH* release into and clearance from the blood. The *LH* release rate is assumed proportional to its amount in the reserve pool, while the clearance rate is proportional to the *LH* blood level.

Likewise, the rate of change of *RP*_*F SH*_ (*t*) in Equation (3) is governed by synthesis and release. The synthesis of *FSH* is inhibited by both *Inh* and *P*_4_. The time delay *τ* is introduced to account for the time it takes the *FSH* synthesis rate to respond to changes in the *Inh* concentration. *P*_4_ stimulates the *FSH* release into the bloodstream. The quadratic expression 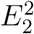 in the *FSH* release term is included to ensure greater inhibitory effect of *E*_2_ on *FSH* release than on *LH* release. Similarly, the rate of change of *FSH*(*t*) in Equation (4) depends on two terms: a release term assumed proportional to the amount of *FSH* in the reserve pool and a clearance term assumed proportional to the *FSH* blood concentration.

Applying this model to the hormone cascade described above, gives the following system of DDEs

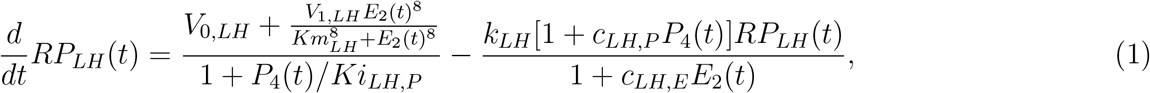

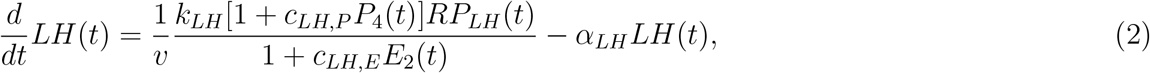

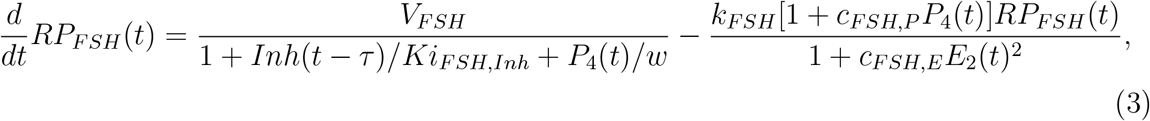

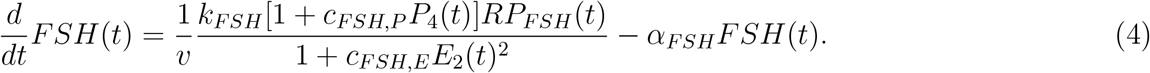

**The ovarian model** predicts the response of the ovarian hormones *E*_2_, *P*_4_, and *Inh* as functions of *LH* and *FSH*.

The rate of change of the mass of active follicular/luteal tissue in Equations (5) to (13) depend on the mass of follicular/luteal tissue promoted to that stage and the mass advancing to the next stage. *FSH* in Equation (5) stimulates, while *P*_4_ inhibits, the recruitment of immature follicles to the *RcF* stage. *LH* in Equations (5) to (7) aids the growth and transition of follicles to the next follicular stage until ovulation.

The rate of production of hormones at each ovarian stage is assumed proportional to the active mass of the follicle or corpus luteum at that stage. Furthermore, the blood concentrations of the ovarian hormones are assumed at a quasi-steady state because the clearance of ovarian hormones from the blood is faster than the clearance of pituitary hormones and the time scale for follicular and luteal development. Thus, the blood concentration of each ovarian hormone in Equations (14) to (16) is written as linear combinations of follicular/luteal masses in the stages secreting it [5, 11, 12, 17]. In addition, exogenous estrogen 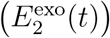 and progesterone 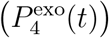, given as blood concentrations, are assumed to affect the body in the same manner as endogenous ovarian hormones. Hence, 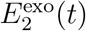 and 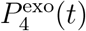 are included in Equations (14) and (15).

Equations (5) to (16) reflect the transition of follicular/luteal mass from recruitment, to ovulation, to the stages of the luteal phase, and the effect of the pituitary to the ovarian hormones, given by the following ordinary differential equations

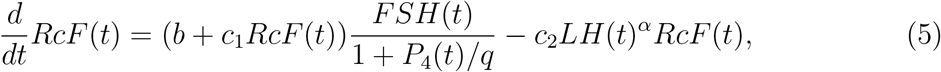

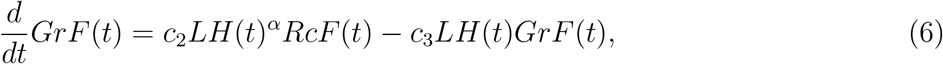

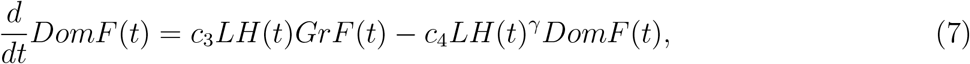

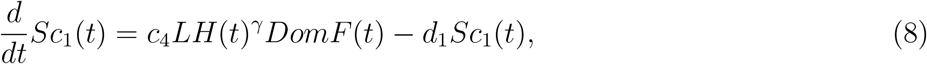

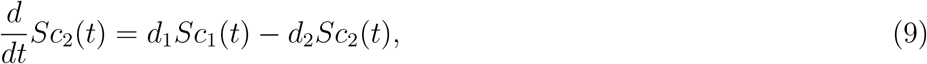

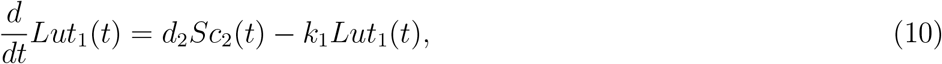

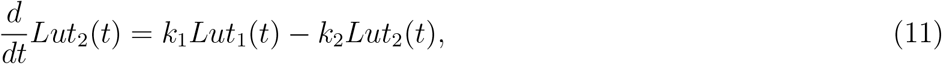

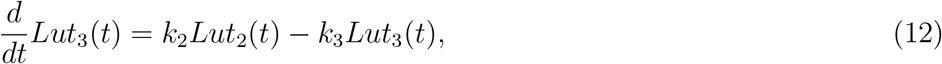

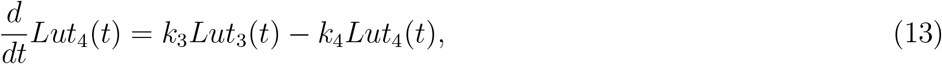

with auxiliary equations

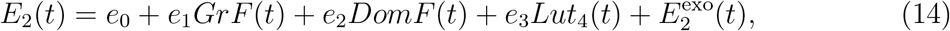

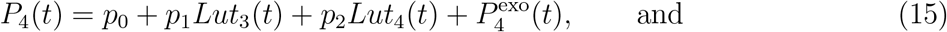

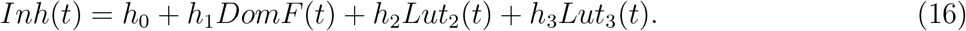

### Novel model modifications: anovulatory cycle and contraception

Exogenous estrogen and/or progesterone inhibit ovulation through different mechanisms. Estrogen causes anovulation by suppressing gonadotropin secretion, depicted in the release term of Equations (1) and (3). Low gonadotropin levels inhibit maturation of follicles causing low production of estrogen insufficient to induce an *LH* surge. The administration of progesterone reduces ovarian hormone levels [30]. The Margolskee model [11] is unable to reflect this condition. Therefore, to attain contraceptive effect of progesterone we included the following terms accounting for

b.1 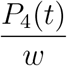 in the synthesis term of Equation (3) and
b.2 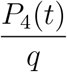 in the first term of Equation (5).

The term 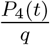 describes the direct inhibitory action of progesterone on follicular development [36, 39]. Baird et al. [36] and Setty et al. [39] also suggested that reduced follicle growth is further caused by the suppression of gonadotropin secretion. Moreover, Batra et al. [40] found that in ovine pituitary cell culture, progesterone decreases *FSH* secretion through decreased *FSH* biosynthesis. This inhibition of *FSH* production is accounted for by the term 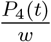. In the model, the administration of both 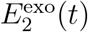 and 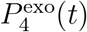 increases suppression of gonadotropin secretion. This leads to lower combined doses to induce anovulation, showing the enhanced effectivity of combined treatment suggested in Rivera et al. [34].

### Parameter estimation

To calibrate the model to the data extracted from Welt et al. [15], we estimated selected model parameters as follows:

c.1 The parameters in the pituitary model Equations (1), (2), (3) (without *P*_4_(*t*)*/w*), and (4) are estimated starting from parameter values and initial conditions in [11]. For this submodel, we replaced *E*_2_(*t*), *P*_4_(*t*), and *Inh*(*t*) by time-dependent functions fitted to *E*_2_, *P*_4_, and *Inh* levels in the data extracted from Welt et al. [15].
c.2 The parameters in the ovarian model Equations (5) (without *P*_4_(*t*)*/q*) to (13), (14) (with 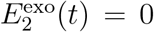), (15) (with 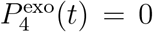), and (16) are estimated starting with parameter values and initial conditions in [11]. For this submodel, we substituted *LH*(*t*) and *FSH*(*t*) by time-dependent functions fitted to *LH* and *FSH* levels in data extracted from Welt et al. [15].
c.3 The parameters obtained in c.1 and c.2 are used to estimate the parameters in the merged pituitary and ovarian model.
c.4 We do not have physiological knowledge of the parameters *w* and *q* thus, values assigned to *w* and *q* with order of magnitude between 0 and 1, and the parameter set obtained in c.3 are used as initial guess to estimate parameters in the final merged model.

In c.1 to c.4 we used the MATLAB `fminsearch` function, which utilizes the Nelder-Mead simplex algorithm, to estimate parameters that minimize least squares error. The least squares function employed in c.4 is

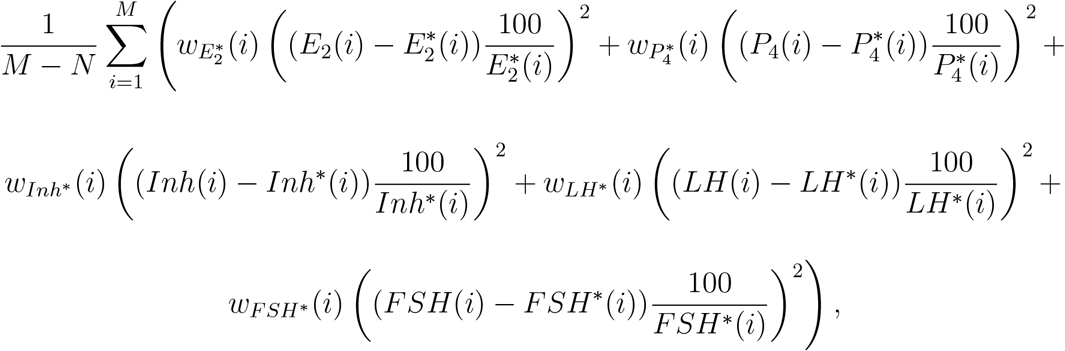

where *H*(*i*) is the hormone model output at day *i, H*^∗^(*i*) is the hormone Welt data at day *i* for hormones *E*_2_, *P*_4_, *Inh, LH*, and *FSH, w*_*H*∗_ (*i*) = *z-score of H*(i)* if *H*(i)* is the maximum of *H\** else, *w*_*H*∗_ (*i*) = 1.

The denominator *M* − *N* is the difference between *M* data points (obtained by repeating the 28-day Welt data four times) and the *N* model parameters and initial conditions, ascertains that errors don’t increase with repetition of the data set. Because the hormones have different units, the multiplier 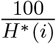 is placed to make the hormones comparable. The *z* − *score*, denoting the number of standard deviations a data point is from the mean, aids in approximating the maximum hormone data. Manual adjustments after optimization are carried out to reach the maximum *E*_2_, *P*_4_, *Inh, LH*, and *FSH*, and to obtain cycle length close to the data cycle length of 28 days. This is essential since an anovulatory cycle is determined by the decrease in *P*_4_ and *LH* maximum levels from the values in a normal cycle. The resulting initial conditions and parameter values are shown in Table I and Table V.

### Optimal control applied to the menstrual cycle model

Optimal control theory describes control strategies to steer a system towards an optimal outcome specified in a cost function [41, 42]. In an optimal control problem, state variables *x*(*t*) depending on time *t*, model a dynamical system and appropriate control function *u*(*t*) is obtained by optimizing an objective function *J* (*u*) subject to constraints [42]. In particular, consider an optimal control problem which minimizes an objective function *J* (*u*) to determine optimal *u*(*t*) and corresponding *x*(*t*). Mathematically, it can be written as

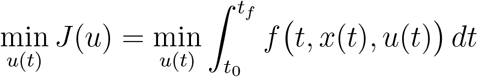

subject to *x*′(*t*) = *g t, x*(*t*), *u*(*t*) and *x*(*t*_0_) = *x*_0_.

In this study, we let 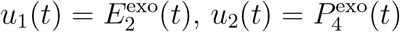, and

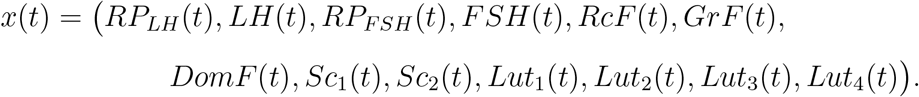

The objective is to seek minimum dosage for *u*_1_(*t*) and *u*_2_(*t*) which decreases the *P*_4_ peak to a value resulting in an anovulatory state. That is,

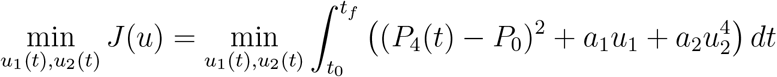

subject to the model *x*′ (*t*) = *g t, t* − *τ, x*(*t*), *u*_1_(*t*), *u*_2_(*t*) and *x*(*t*_0_) = *x*_0_.

In the integrand, the target point *P*_0_ = 4 ng/mL per day in the first term is chosen to keep *P*_4_(*t*) *<* 5 ng/mL for 0 = *t*_0_ ≤ *t* ≤ *t*_*f*_ = 28. The second and third terms are included to minimize the dosage of exogenous hormones. The power of *u*_2_ is 4 in order to add convexity to the optimal control problem and to smoothen the control. Very oscillatory results are obtained for power of *u*_2_ less than four. To further smoothen results other methods may be explored like varying initial conditions and number of time points. The constants *a*_1_ *>* 0 and *a*_2_ *>* 0 are weight factors which balance the effort of minimizing *u*_1_ and *u*_2_, and hitting *P*_0_. These weights are selected using an iterative process to produce desired *P*_4_ levels.

To solve the optimal control problem we applied control parameterization. In this numerical method the control function is approximated by a linear combination of basis functions [41, 43]. Specifically, we utilize the MATLAB `makima` function which performs a piecewise cubic Hermite interpolation to estimate the control function. Moreover, we use `dde23` to solve the delay differential equations and `fmincon` to generate components of the piecewise cubic polynomial which gives the local least cost.

### Exploring effects of exogenous hormones

To investigate the effect of exogenous estrogen and progesterone on the menstrual cycle, we compared the

d.1 model output without exogenous hormone to data extracted from Welt et al. [15] for normally cycling women repeated four times,
d.2 model output with constant dose of exogenous hormone to hormone output in d.1, and
d.3 model output with optimal time-varying dose of exogenous hormone to hormone outputs in d.1 and d.2.

Some types of hormonal contraceptives like implants, injections, and patches are administered non-orally and continuously [21] while birth control pills are taken orally at specific time points. In d.2 and d.3 we study model response to exogenous estrogen and progesterone monotherapy, and combined treatment administered continuously for one cycle (28 days).

## RESULTS

This section presents the model output with and without administration of exogenous hormones. Sum of squared residuals between the model output without administration of exogenous hormones and Welt data, model output peak, and period are shown. In the administration of exogenous hormones, constant and time-varying doses are considered. Exogenous estrogen and progesterone monotherapies, and combination treatment of the two hormones are explored. For each type of therapy, minimum dose of exogenous estrogen and/or progesterone which leads to anovulation is determined. Optimal control theory is applied to investigate optimal time-varying doses.

### The normal cycle solution

Without the administration of exogenous hormones (i.e., 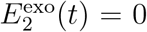 and 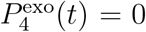), the estimated initial condition in Table I produces a unique stable periodic solution (called the normal cycle solution) to the menstrual cycle model. The cycle length is 28.05 days. Local stability of the solution is affirmed by varying the initial conditions.

Figure 4 depicts the dynamics of the ovarian and pituitary hormones *E*_2_, *P*_4_, *Inh, LH*, and *FSH* predicted by the model. The normal cycle solution (blue curve) exhibits hormone surges and dips and is a good estimate to the data extracted from Welt et al. [15] (red curve). Table II presents the sum of squared residuals between the model output and Welt data, and hormone output peaks.

**TABLE II.** Sum of squared residuals between the model output and Welt data, and model output peaks.

| Hormone |  | Sum of squared residuals |
| --- | --- | --- |
| $E_2$ (in $\text{pg}^2/\text{mL}^2$) | | 21839.63 |
| $P_4$ (in $\text{ng}^2/\text{mL}^2$) | | 182.77 |
| $Inh$ (in $\text{IU}^2/\text{mL}^2$) | | 66.55 |
| $LH$ (in $\text{IU}^2/\text{L}^2$) | | 1579.46 |
| $FSH$ (in $\text{IU}^2/\text{L}^2$) | | 174.19 |
| Hormone peak |  |  |
| Hormone | Welt data | Model output |
| $E_2$ (in $\text{pg}/\text{mL}$) | 237 | 237.74 |
| $P_4$ (in $\text{ng}/\text{mL}$) | 17.9 | 18.06 |
| $Inh$ (in $\text{IU}/\text{mL}$) | 11.5 | 11.47 |
| $LH$ (in $\text{IU}/\text{L}$) | 123 | 123.12 |
| $FSH$ (in $\text{IU}/\text{L}$) | 19.6 | 20.36 |

**FIG. 4.**
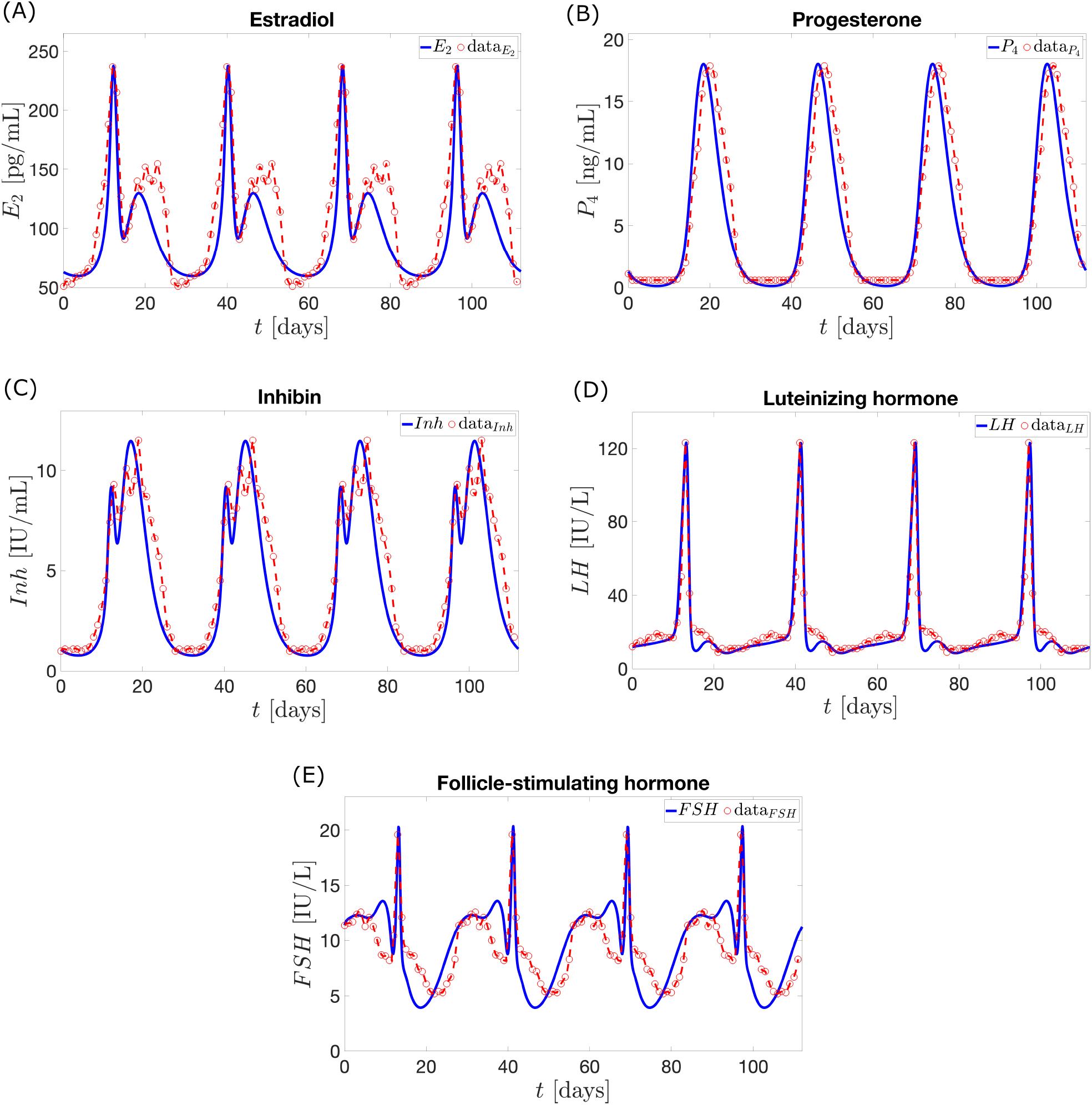
The blue curves describe the dynamics of the pituitary and ovarian hormones predicted by the model without exogenous estrogen and progesterone. The red curves represent the 28-day normal cycle hormone data extracted from Welt et al. [15] repeated for 112 days. The model output, which has period 28.05 days, gives a good fit as shown in Table II.

### Administration of exogenous hormones

#### Constant dosage

Exogenous estrogen and/or progesterone inhibits pituitary and ovarian maximum hormone levels [30, 31]. To simulate the response to a constant dose of exogenous estrogen monotherapy, 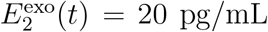 per day and 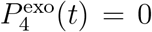 are used in Equation (14) and Equation (15) for 28 days. Similarly, the effect of a constant dose of exogenous progesterone monotherapy is obtained with 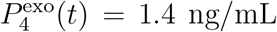 per day and 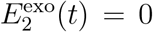. Table III presents the percentage decrease in model output peak compared to the hormone output peak of the Wright model of hormonal contraception [13]. The use of 20 pg/mL per day of estrogen or 1.4 ng/mL per day of progesterone results to the hormone profiles in Figure 5. Each of these amounts is insufficient to manifest anovulation because although reduced, the maximum *P*_4_ value is still more than 5 ng/ml. To determine 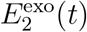 and 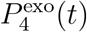 values which block ovulation, we observe the *LH* and *P*_4_ model output as the dosages vary from 0 to 60 pg/mL per day for 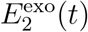, and 0 to 4 ng/mL per day for 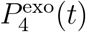. Figures 6 and 7 illustrate that increasing dosage leads to eliminating *LH* surge and decreasing fluctuation in *P*_4_ level. In the estrogen monotherapy, higher 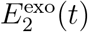 generates lower maximum *P*_4_ and anovulation is attained when 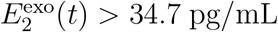. In progesterone monotherapy, ovulation is suppressed between doses 3.1 ng/mL and 3.7 ng/mL. The small window of anovulation is due to 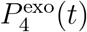 in Equation (15), and the linear inhibitory terms *P*_4_(*t*)*/w* and *P*_4_(*t*)*/q* in Equation (3) and Equation (5). Because the inhibitory term *P*_4_(*t*)*/w* is linear, to bring *P*_4_(*t*) *<* 5 ng/mL the amount of *P*_4_(*t*) must provide strong inhibition of *FSH*. This is attained by increasing 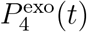 in Equation (15). A high amount of 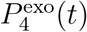 also enhances inhibition of the linear term *P*_4_(*t*)*/q* on *RcF* (*t*), which in turn suppresses further *Lut*_3_(*t*) and *Lut*_4_(*t*). However, because of the inclusion of 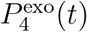 in Equation (15), increasing amount of 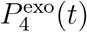 also results to raising *P*_4_(*t*) toward values more than 5 ng/mL, which are ovulatory levels.

**TABLE III.** Percentage decrease in model output peak with the administration of exogenous estrogen/progesterone.

| Model output peak percentage decrease |  |  |
| --- | --- | --- |
| $E_2^{\text{exo}}(t) = 20 \text{ pg/mL per day}$ | Current model | Wright model |
| $E_2$ (in pg/mL) | 21 | 16 |
| $P_4$ (in ng/mL) | 35 | 23 |
| $Inh$ (in IU/mL) | 33 | 22 |
| $LH$ (in IU/L) | 12 | 26 |
| $FSH$ (in IU/L) | 9 | 13 |
| $P_4^{\text{exo}}(t) = 1.4 \text{ ng/mL per day}$ | Current model | Wright model |
| $E_2$ (in pg/mL) | 51 | 81 |
| $P_4$ (in ng/mL) | 53 | 81 |
| $Inh$ (in IU/mL) | 55 | 82 |
| $LH$ (in IU/L) | 81 | 89 |
| $FSH$ (in IU/L) | 35 | 29 |

**FIG. 5.**
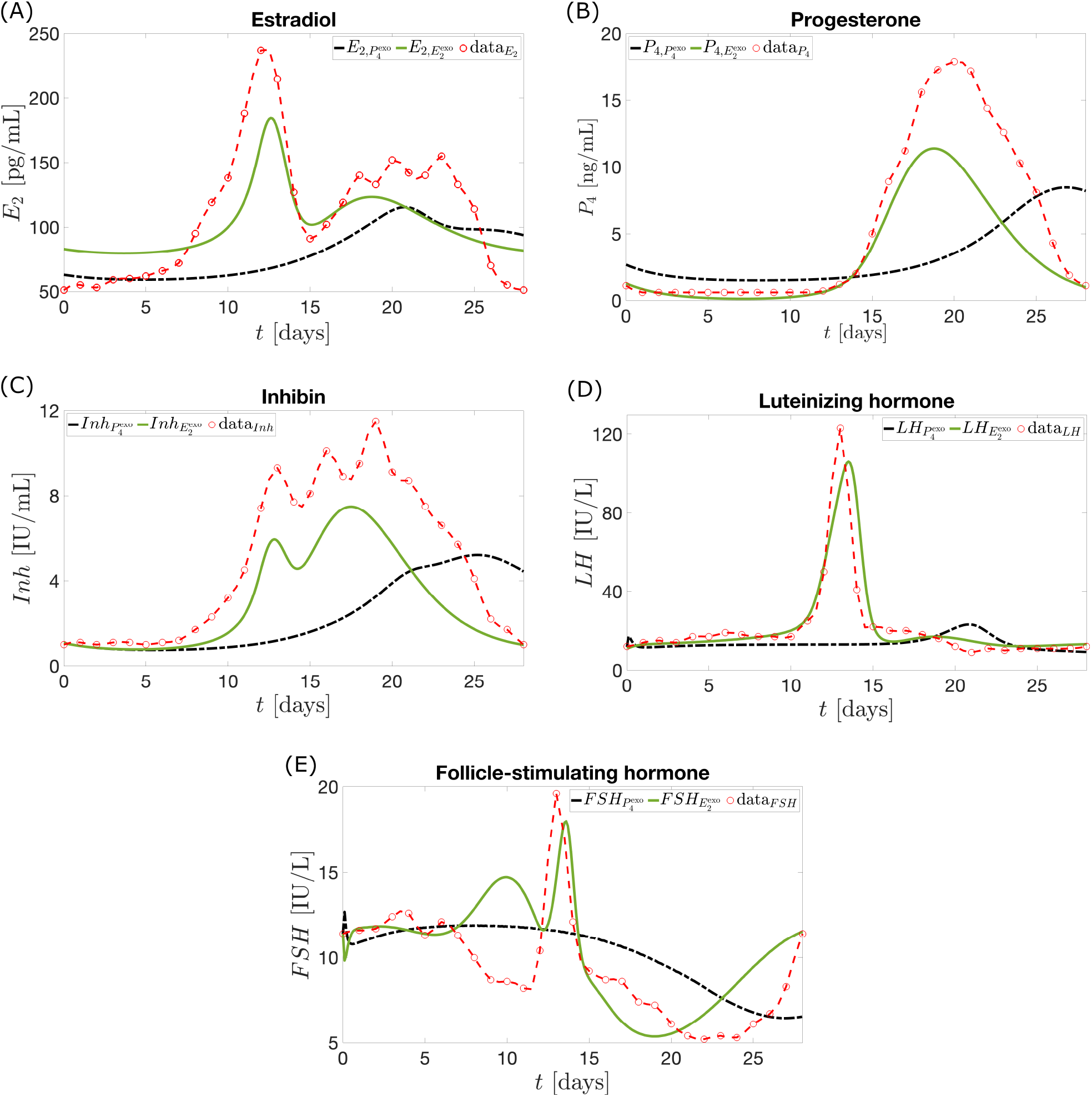
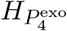 (black curve) or 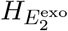 (full green curve) is the hormone *H* model output in case of 20 pg/mL per day of exogenous *E*_2_ or 1.4 ng/mL per day of exogenous *P*_4_ is administered for 28 days, respectively. The stipulated red curve represents the data for the 28-day normal cycle extracted from Welt et al. [15]. The addition of exogenous *E*_2_ or *P*_4_ reduces the peak of each of the five hormones.

**FIG. 6.**
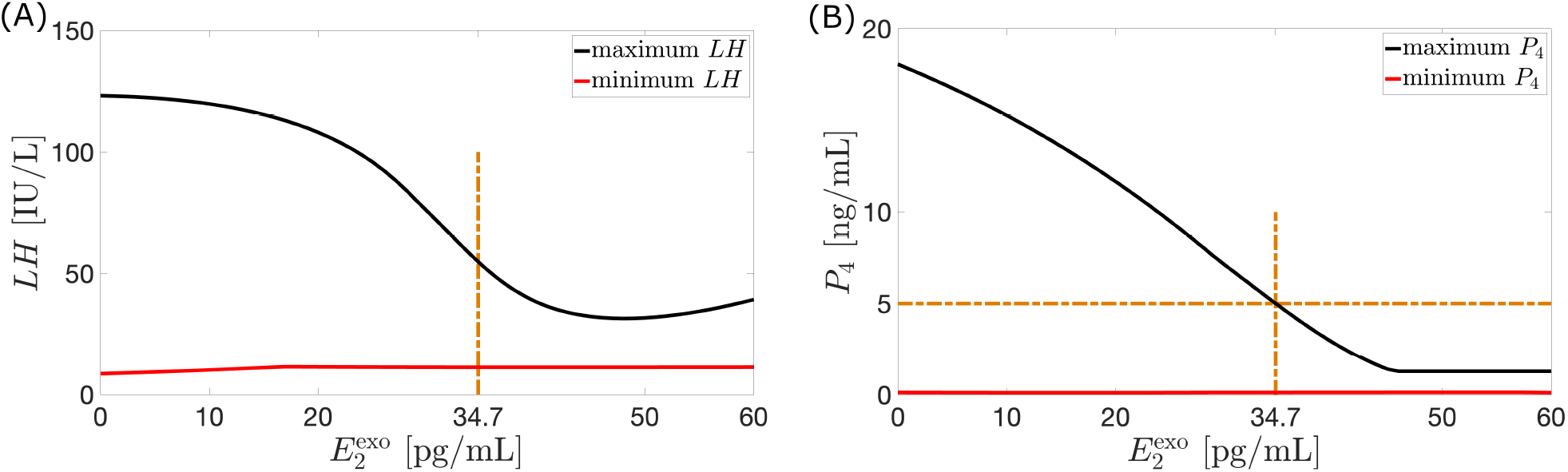
The vertical axes in (A) and (B) present the maximum (full black curve) and minimum (full red curve) values over a 28-day cycle reached by *LH* and *P*_4_, respectively, when the corresponding amount of exogenous *E*_2_ is given. Panel (B) is similar to (A) but shows that increasing dosage of 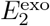 causes decreasing amplitude of the variation in *P*_4_ level. Anovulation is attained when 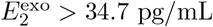.

**FIG. 7.**
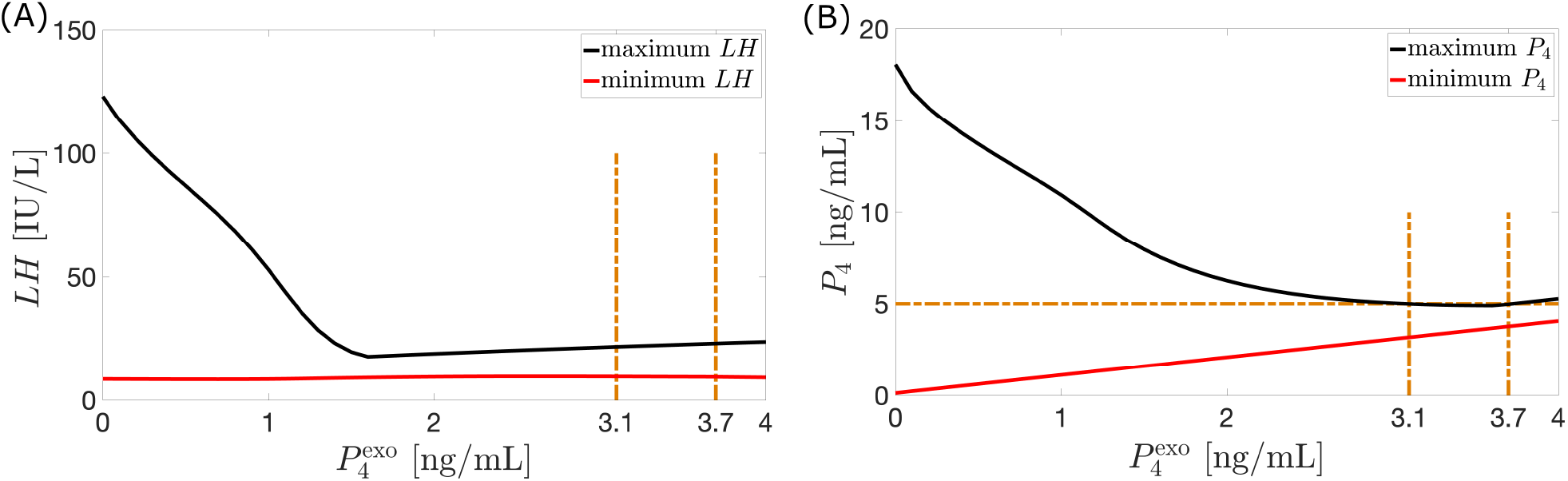
Shown in (A) and (B) are the maximum (full black curve) and minimum (full red curve) values attained by *LH* and *P*_4_ over a 28-day cycle resulting from the administration of the corresponding dosage of 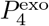. Panel (B) illustrates diminishing fluctuation in *P*_4_ value. Anovulation is achieved between 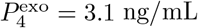 and 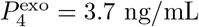.

As in [13], an anovulatory effect of a high dosage of exogenous estrogen or progesterone can be achieved by using a combination of the two hormones. For instance, monotreatment by 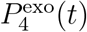 between 3.1 ng/mL per day and 3.7 ng/mL per day or 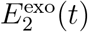 greater than 34.7 pg/mL per day for the entire 28-day cycle induces anovulation see Figures (6(B) and 7(B)). Figure 8(A) shows that alternatively, anovulation can be obtained by a combination treatment with 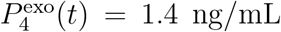 per day and 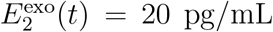 per day. Recall that the monotreatment with 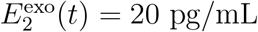 per day or 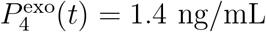 per day does not prevent ovulation see Figure 5(B). Other combination treatments which block ovulation are depicted in Figure 8(B).

**FIG. 8.**
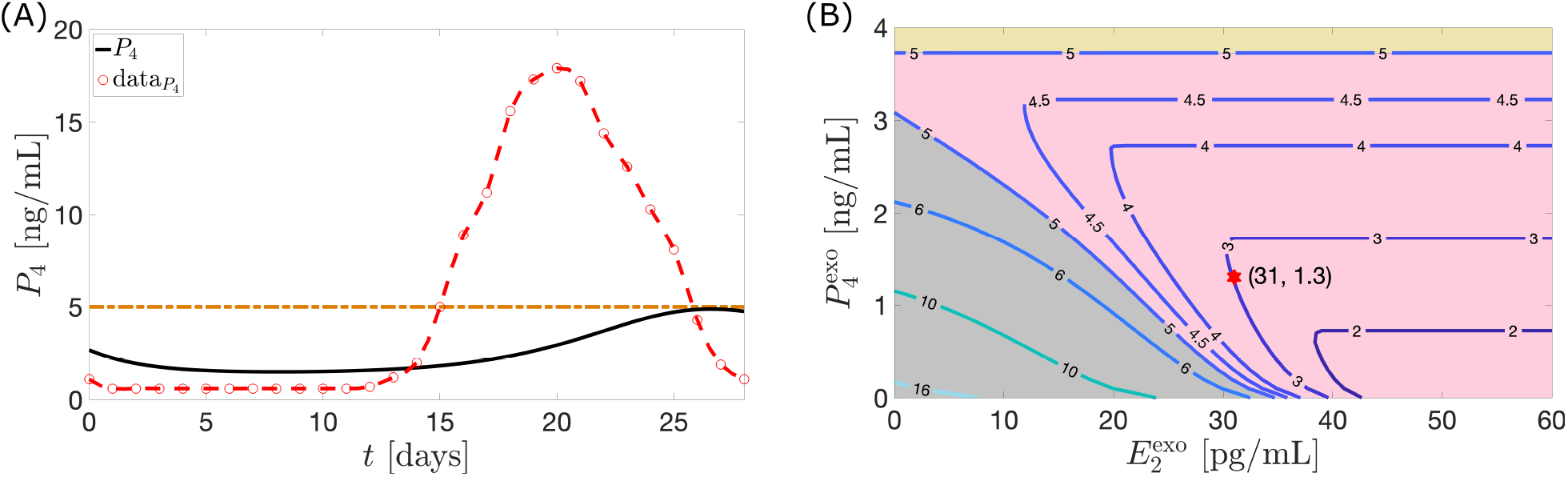
(A) The black curve represents the *P*_4_ model output for a combination treatment with 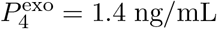 per day and 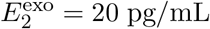 per day administered for 28 days. The red circles with the interpolated stipulated curve represents the 28-day normal cycle hormone data extracted from Welt et al. [15]. The treatment suppresses the *P*_4_ concentration achieving anovulatory state. In (B) the curves composed of points 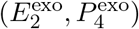, correspond to a combined dosage of 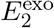 and 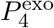 resulting in a *P*_4_ maximum value of *k* ng/mL. Anovulation is attained when *k <* 5. The yellow region 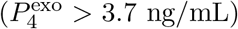 corresponds to ovulation. On the lower left portion, an almost straight line with slope −0.1 separates the regions of ovulation (in gray) and anovulation (in pink).

Figure 8(B) shows contour plots of maximum *P*_4_ levels over a cycle for various combination treatments of 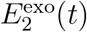 and 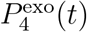. For instance, the point (31, 1.3) on the contour labeled 3 signifies that an infusion of a combination of 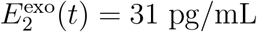 per day and 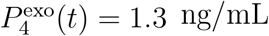 per day yields a maximum *P*_4_ level of 3 ng/mL. The region of anovulation (in pink) is bounded above and on the left by the curves *k* = 5. The left boundary is approximately a straight line with slope −0.1. Thus, to stay on this boundary a reduction of 1 pg/mL per day of 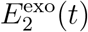 needs to be counteracted by an increase of approximately 0.1 ng/mL per day of 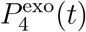.

### Optimal nonconstant dosage

This section uses optimal control to determine optimal time-varying doses that induce anovulation. In order to explore mono and combination treatments, three different cases of the objective function are examined.

#### Case 1. Exogenous estrogen as monotherapy

Let *u*_2_(*t*) = 0. For *a*_1_ = 0.4 *µ*g/mL then the control *u*_1_(*t*) which minimizes the objective function is illustrated in Figure 9(A).

**FIG. 9.**
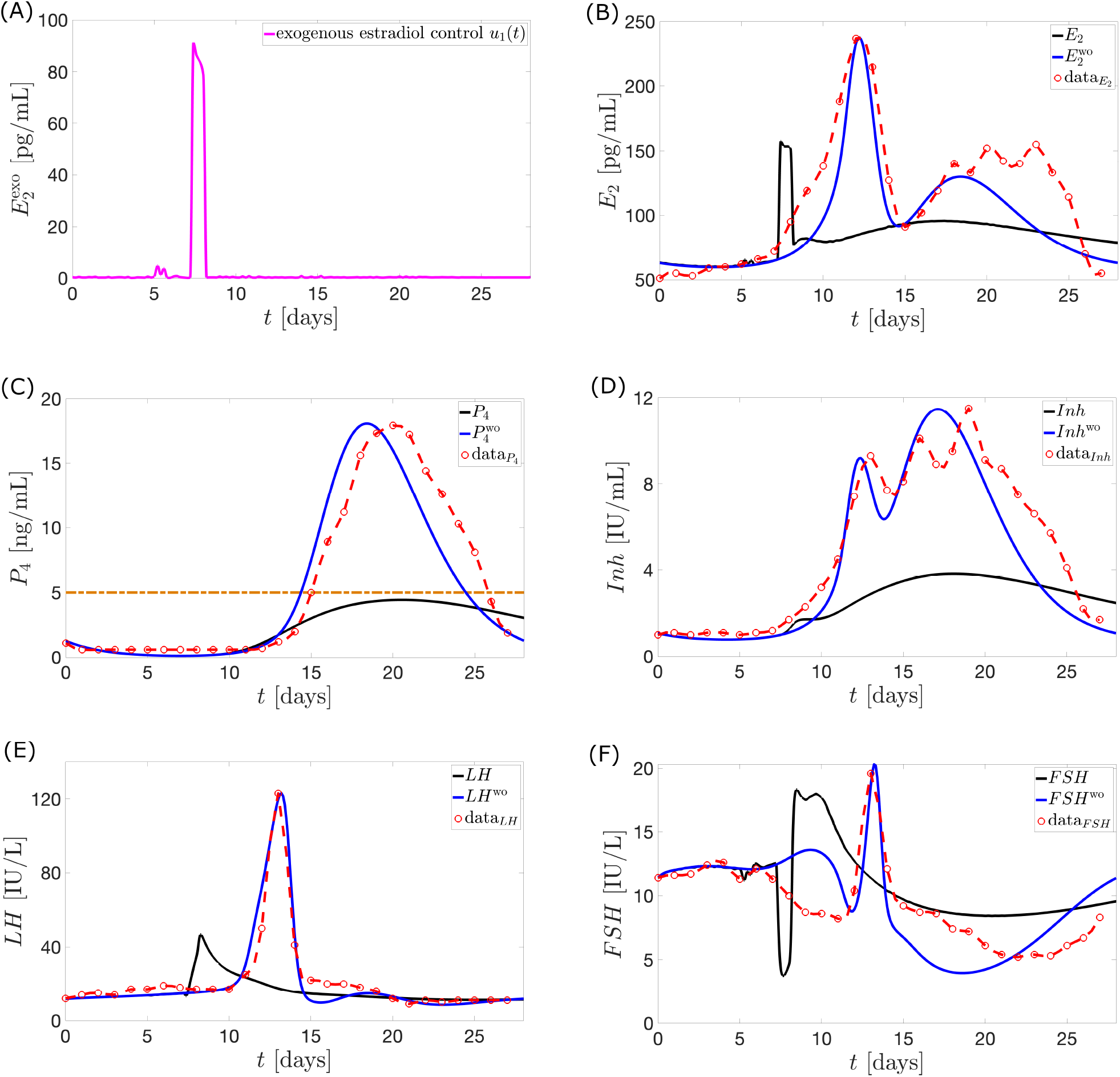
The full black curve is the model output when the optimal control *u*_1_ (magenta in panel (A)) is applied. *H*^wo^ (full blue curve) denotes the hormone model output without the influence of *u*_1_. This is the normal cycle solution. The Welt data for a normal cycle is presented in red circles with the interpolated curve. The maximum *P*_4_ value is 4.43 ng/mL.

In Figure 9, the steep rise in the dosage of exogenous estrogen in the optimal control inhibits strongly the release of *FSH* in the bloodstream via Equation (3). This causes the *FSH* levels around the time of *u*_1_ surge to plunge. Consequently, a low mass of *Rc*F (see Figure 10) is produced via Equation (5). The underdeveloped follicles then produce lower *E*_2_. The lower concentration of *E*_2_ causes a decrease in *LH* production via Equation (1). Without an *LH* surge, ovulation does not occur. This implies the value of *P*_4_ to be lowered via Equation (14).

**FIG. 10.**
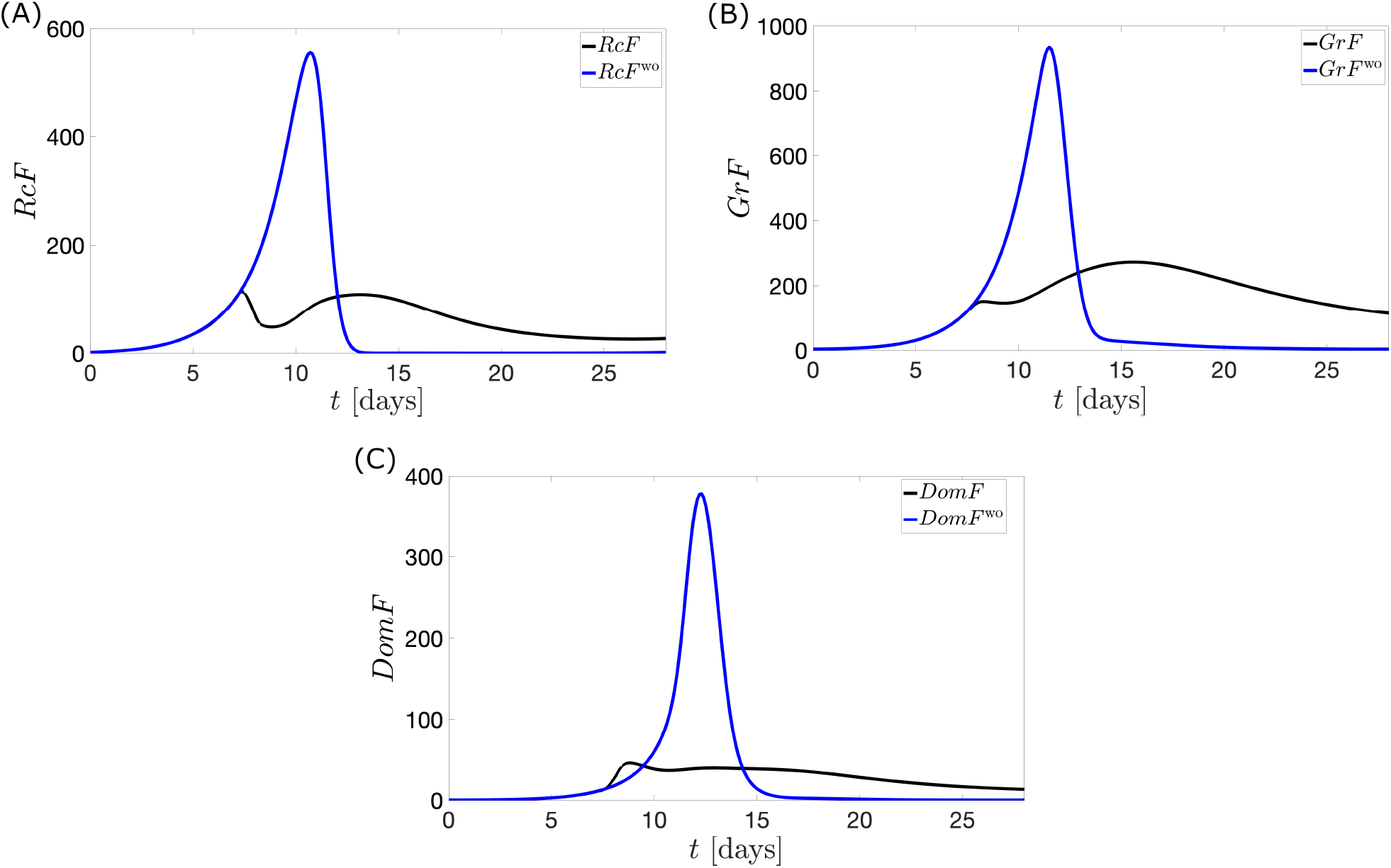
The full black curve describes the follicular mass when the optimal control *u*_1_ is applied. The full blue curve shows the follicular mass without the application of *u*_1_. In (A), the steep decline in *RcF* is evident on the interval of *FSH* level drop in Figure 9(F). The inhibition of *RcF* subsequently contributes to the reduced development of *GrF* and *DomF* in panels (B) and (C).

The large dosage of 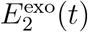 given in the mid-follicular phase is effective in preventing the ovulatory level of *E*_2_ in the late follicular phase. The schedule of administration agrees with [44] that estrogen treatment started before the 10th day of the menstrual cycle can result in an anovulation.

Increasing the value of *a*_1_ penalizes *u*_1_ dose more strongly. This reduces *u*_1_ dose further but increases deviation of *P*_4_ from 4 ng/mL.

#### Case 2. Exogenous progesterone as monotherapy

Assume *u*_1_(*t*) = 0 and *a*_2_ = 0.07 mL^2^/ng^2^. Anovulatory hormonal levels result from a continuous suppression of *FSH* levels (see Figure 11(F)), in contrast to the sudden dip in Figure 9(F).

**FIG. 11.**
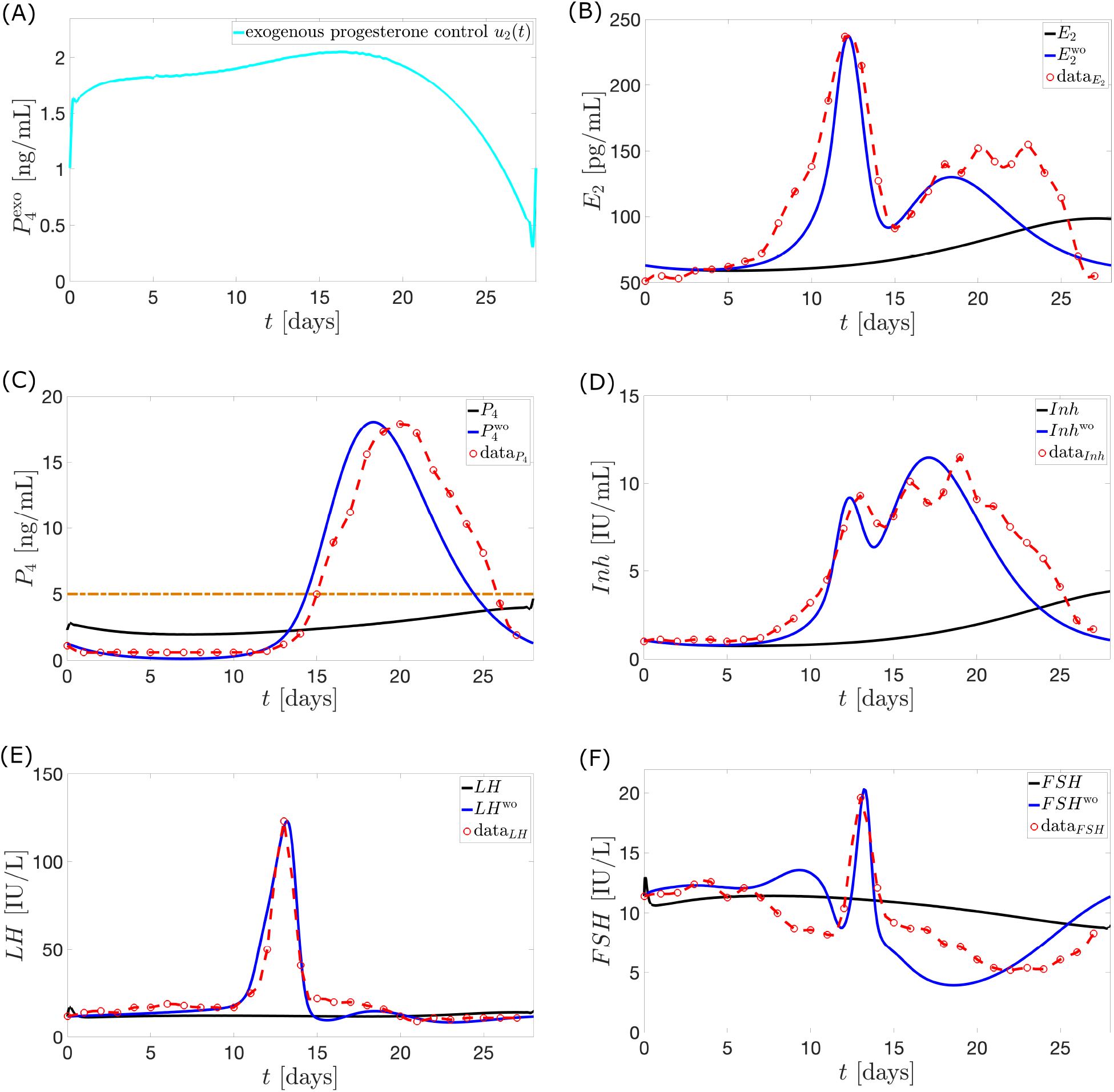
The full black curve is the model output when the optimal control *u*_2_ (cyan in panel (A)) is used. The normal cycle solution *H*^wo^ (full blue curve) is the hormone model output when *u*_2_ is not administered. The red circles with the interpolated curve denote the Welt data for a normal cycle. *P*_4_ reaches a maximum value of approximately 4.66 ng/mL.

The administration of *u*_2_ starting from the first day of the cycle does not permit *FSH* to reach its maximum value due to the low synthesis in the pituitary via Equation (3). The low level of *FSH* in the follicular phase and the additional inhibition by *P*_4_ via Equation (5) hinder follicular growth. Consequently, *E*_2_ levels far lower than normal are attained via Equation (14). Nonoccurrence of *LH* surge follow via Equation (1).

The optimal control *u*_2_ suggests that a maximum dosage be given before the time in the normal menstrual cycle when *P*_4_ peaks. *Inh* prevents *FSH* synthesis via Equation (3) but anovulation results to decreased *Inh* levels via Equation (16). Thus, the mathematical model causes *u*_2_ to still have high doses after day 14 in order to continue suppression of *FSH* production by compensating for the reduced inhibition by *Inh*.

Similar to Case 1, increasing value of *a*_2_ puts more effort in minimizing *u*_2_ dose, increasing deviation of *P*_4_ from 4 ng/mL.

#### Case 3. Administration of combined exogenous estrogen and progesterone

Allowing *u*_1_(*t*) ≠ 0, *u*_2_(*t*) ≠ 0, *a*_1_ = 0.4 *µ*g/mL and *a*_2_ = 0.7 mL^2^/ng^2^, then the optimal controls *u*_1_ and *u*_2_ illustrated in Figure 12(A) yield the dynamics in Figure 12(B) to Figure 12(F).

**FIG. 12.**
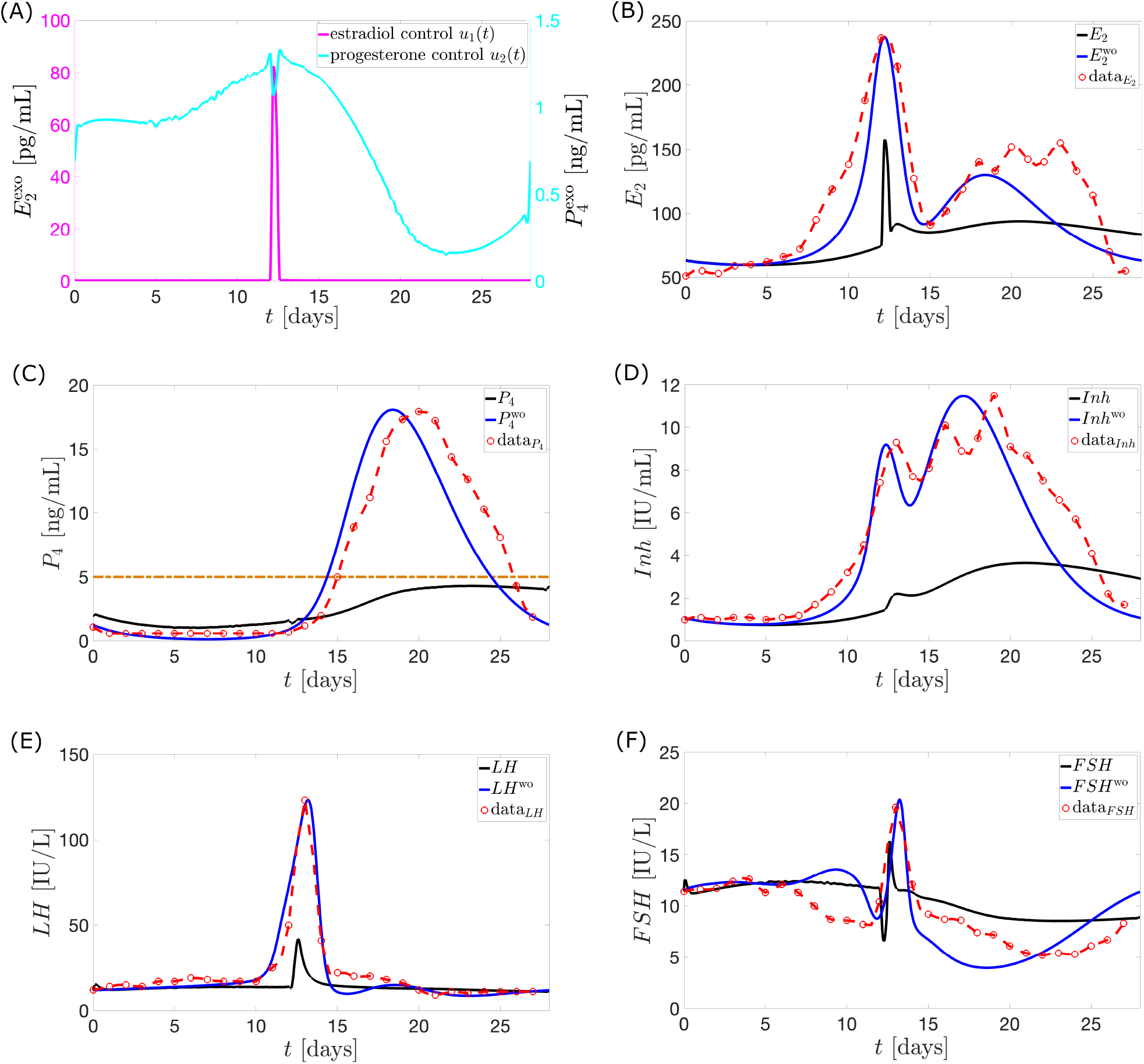
The full black curve is the model output when the optimal controls *u*_1_ and *u*_2_ (magenta and cyan in panel (A)) are administered. The normal cycle solution *H*^wo^ (full blue curve) is the hormone model output when *u*_1_ and *u*_2_ are not applied. The red circles with the interpolated curve denote the Welt data for a normal cycle. The maximum *P*_4_ concentration is approximately 4.31 ng/mL.

The weights attached to the *u*_1_ and *u*_2_ terms result to smaller percentage decrease of *u*_1_ peak from Case 1 compared to the percentage decrease of *u*_2_ peak from Case 2. This leads to a greater influence of the optimal control *u*_1_ on the menstrual cycle. Figure 12 presents hormone profiles resembling the ones shown in Figure 9. The weight *a*_2_ may be decreased if the intention is to diminish the impact of *E*_2_.

Not only does the combination therapy utilize lower doses of exogenous estrogen and progesterone, it also allows the administration of *u*_1_ to commence in a later follicular stage. The late administration of *u*_1_ is possibly compensated by the inhibition provided by exogenous *P*_4_ in the early follicular phase. The surge in the *u*_1_ value around *t* = 12.2 days allowed a dip in the *u*_2_ amount. The combination of the *u*_1_ peak dose and the *u*_2_ dose at this time is able to maintain anovulation. Unlike in Case 1, the dosage and timing of administration of *u*_1_ does not suffice to keep anovulatory *P*_4_ level until the end of cycle thus, use of increasing *u*_2_ is needed in the late luteal phase.

## DISCUSSION

With the rapid development of new implants and injections providing continuous administration there is great potential to implement new treatment scheme minimizing dose. This study employs optimal control to a modification of the model in Margolskee et al. [11] to determine the optimal time-varying dose of exogenous estrogen and/or progesterone that induce anovulation. The parameter estimation employed yields a model output that predicts well the data extracted from Welt et al. [15]. An improvement to previous mathematical models, hormone output peaks are close to the data peaks (see Table II). This is essential since an anovulatory cycle is determined from the reduction of the normal maximum *P*_4_ level to less than 5 ng/mL and the lack of *LH* surge. The cycle length is also close to the 28-day period of the Welt data. This makes the model beneficial in future studies investigating the effect of exogenous hormones on cycle length.

Exogenous estrogen and/or progesterone inhibits pituitary and ovarian maximum hormone levels [30, 31]. In the model, the administration of exogenous estrogen and/or progesterone caused a reduction in maximum hormonal values (see Figure 5). Such effect is also produced by the hormonal contraception model in Wright et al. [13]. With 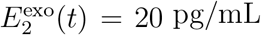 per day, the current model suppresses more the peaks of three of the five hormones but with 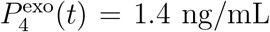 per day, the Wright model reduces more the peaks of four of the five hormones (see Table III). The model in Wright et al. [13] provides greater repression by exogenous *P*_4_ because it uses a nonlinear term to inhibit *RcF* growth and an additional equation depicting upregulation of *P*_4_ by *E*_2_, boosting the contraceptive effect of *P*_4_. We opted for the linear inhibitory term *P*_4_(*t*)*/q* and fewer additions to the Margolskee model [11] to keep the current model simple, reducing the computation time in running our optimization code. This is because the numerical method used in this study, control parameterization using MATLAB functions `dde23` and `fmincon`, though easy to implement is not cost effective.

To our knowledge there has been no study applying optimal control theory on the menstrual cycle model. Optimal control could provide drug administration scheme which greatly enhances contraception outcome by significantly minimizing risks associated with high doses such as venous thromboembolism and myocardial infarction [3, 25–27]. The study by Gu et al. [45] showed that compared to constant-dose administration, optimal control results could substantially improve HIV treatment. Results of our work similarly suggest the significant advantage of optimal time-varying doses.

In estrogen monotherapy, the minimum constant dosage of estrogen over 28 days resulting in an anovulation is (34.73 pg/mL)*×*28 = 972.44 pg/mL (see Figure 13(A)). This dosage lowers the maximum *P*_4_ level to 4.99 ng/mL. The administration of the optimal control *u*_1_ is able to bring down the maximum *P*_4_ level to 4.43 ng/mL (i.e., anovulation is achieved) with only a total dosage (area under the curve or AUC) of 77.76 pg/mL (see Figure 13(B)). A dosage of 894.68 pg/mL (about 92% of minimum total constant dosage) would be saved if *u*_1_ is used to induce anovulation.

**FIG. 13.**
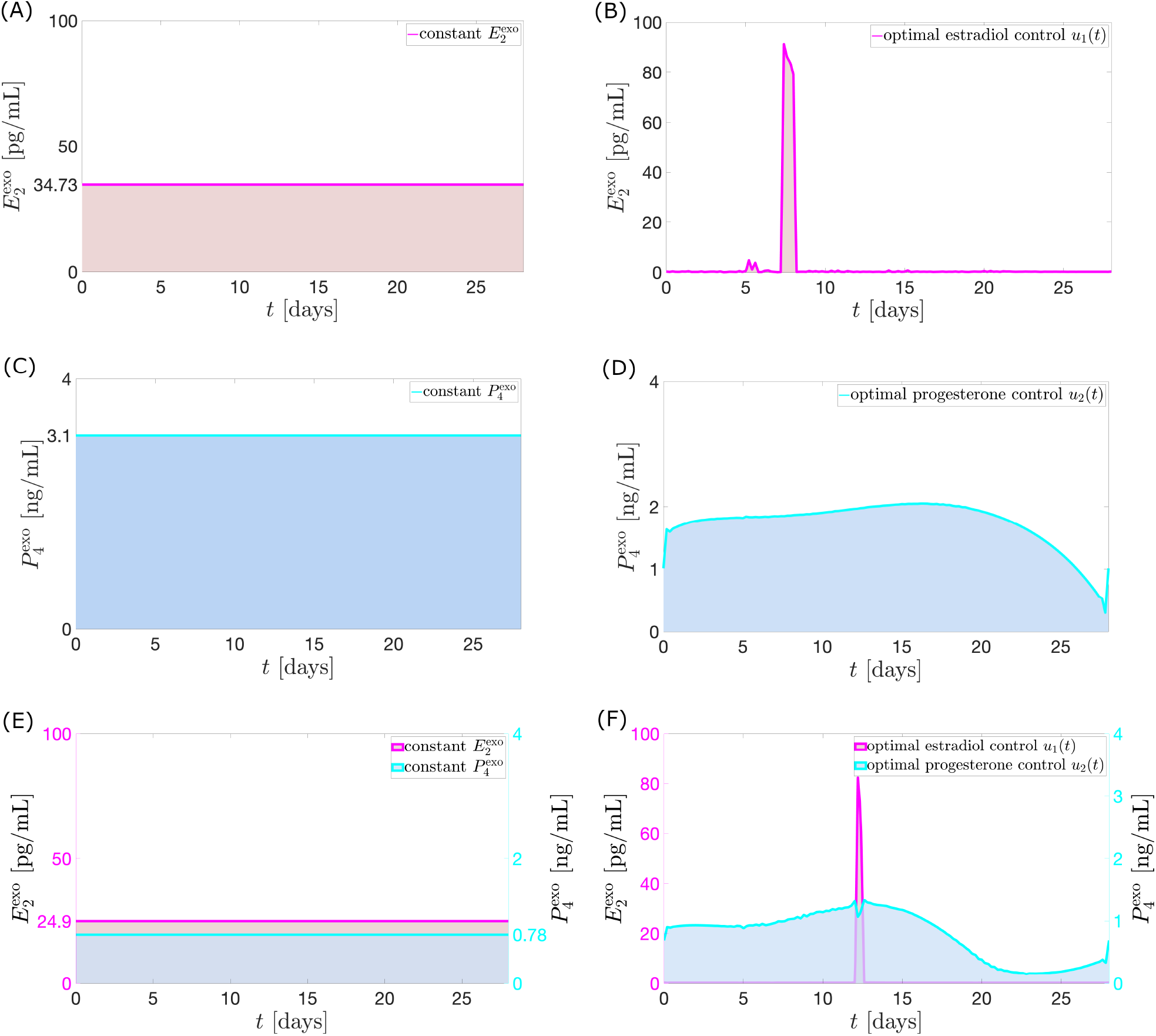
The shaded regions in Panels (A), (C), and (E) indicate the minimum total constant dosage of exogenous estrogen and/or progesterone over 28 days that lowers maximum *P*_4_ concentration to 4.99 ng/mL. The shaded region below *u*_1_ (area under the curve or AUC) in Panel (B) is the total nonconstant dosage of exogenous *E*_2_ which suppresses the *P*_4_ level to 4.43 ng/mL, a reduction by about 92% of the total dosage in (A). Panel (D) illustrates the total nonconstant dosage of exogenous *P*_4_ that reduces maximum *P*_4_ to 4.66 ng/mL, a reduction by about 43% of the total dosage in (C). Panel (F) shows the combined nonconstant doses of exogenous *E*_2_ and *P*_4_ that gives a maximum *P*_4_ level of 4.31 ng/mL.

In the progesterone monotherapy, *u*_2_ is able to bring down the maximum *P*_4_ level in 28 days to 4.66 ng/mL with only a total dosage (AUC) of 48.84 ng/mL (see Figure 13(D)). A constant administration would require 3.1 ng/mL per day, a total dosage of (3.1 ng/mL)*×*28= 86.8 ng/mL to lower the maximum *P*_4_ to 4.99 ng/mL (see Figure 13(C)). A dosage of 37.96 ng/mL (about 43% of the total dosage for constant administration) would be saved if *u*_2_ is employed.

Now in the combination therapy, the total dosage given by *u*_1_ (AUC*u*_1_) is 35.58 pg/mL while that of *u*_2_ (AUC*u*_2_) is 21.67 ng/mL (see Figure 13(F)). If AUC*u*_2_ is taken and spread out constantly in 28 days, then 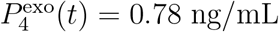 (see Figure 13(E)). Consider the least amount of 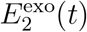 administered constantly in combination with this 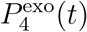 that results to anovulation. Guided by the contour plot (see Figure 8(B)), 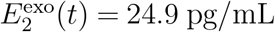 suppresses maximum *P*_4_ to 4.99 ng/mL. The difference between the total exogenous estrogen dosage between the constant and nonconstant administration is (24.9 pg/mL)*×*28 − AUC*u*_1_ = 661.62 pg/mL. Hence about 94.89% of the total 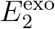 dosage for constant administration would be saved if the combination of *u*_1_ and *u*_2_ is taken. On the other hand, if the total dosage (AUC*u*_1_) given by *u*_1_ is spread out constantly in 28 days, then 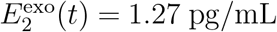. The contour plot (see Figure 8(B)) implies that the least amount of 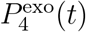 which can be given constantly in combination with 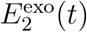 to decrease maximum *P*_4_ to 4.99 ng/mL is 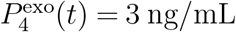. The difference between the total 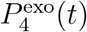 dosage between the constant and nonconstant administration is (3 ng/mL)*×*28 − AUC*u*_2_ = 62.33 ng/mL. Hence about 74.20% of the total 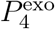 dosage for constant administration would be saved if the combination of *u*_1_ and *u*_2_ is used to inhibit ovulation. A summary of the results of the optimal control strategies is shown in Table IV.

**TABLE IV.** Maximum progesterone level throughout a menstrual cycle caused by the indicated total dose of the optimal exogenous hormone.

| **Treatment regimen** | **Total dose** | **maximum $P_4$ level** |
| --- | --- | --- |
| estrogen monotherapy | 77.76 pg/mL | 4.43 ng/mL |
| progesterone monotherapy | 48.84 ng/mL | 4.66 ng/mL |
| combination therapy | estrogen = 35.58 pg/mL<br>progesterone = 21.67 ng/mL | 4.31 ng/mL |

The optimal control results provide the best timing of administration since an earlier or delayed application of *u*_1_ and/or *u*_2_ yields higher *P*_4_ level. For instance, if *u*_1_ is applied four times in equal intervals from day 0 to day 112 (see Figure 14(A)), anovulation is no longer induced beginning on the third 28-day period (i.e. from day 56 to day 112). This is because the application of *u*_1_ in the preceding period changes the dynamics of the menstrual cycle in the succeeding days. One of these changes is the cycle length. Now, because the control *u*_1_ must be administered at a time before *E*_2_ surge (when *u*_1_ is not applied), the timing of administration in the next cycles must also be changed to continuously suppress ovulation. Sometimes resulting cycle length is less than 28 days so only a portion of *u*_1_ will be applied. In addition, because lower levels of *u*_1_ are negligible compared to the higher levels, we applied only the large-dose portion of *u*_1_ from day 28 to day 112 (see Figure 14(B)). The administration is done when *E*_2_ level is increasing and reaches 75 pg/mL. We are currently conducting further investigation to determine the highest *E*_2_ hormone level where administration of *u*_1_ would commence to still inhibit ovulation. Note that although *u*_1_ is applied six times (in unequal intervals) from day 0 to day 112, the total dosage is still significantly lower than the total constant dosage of exogenous *E*_2_ which induces anovulation.

**FIG. 14.**
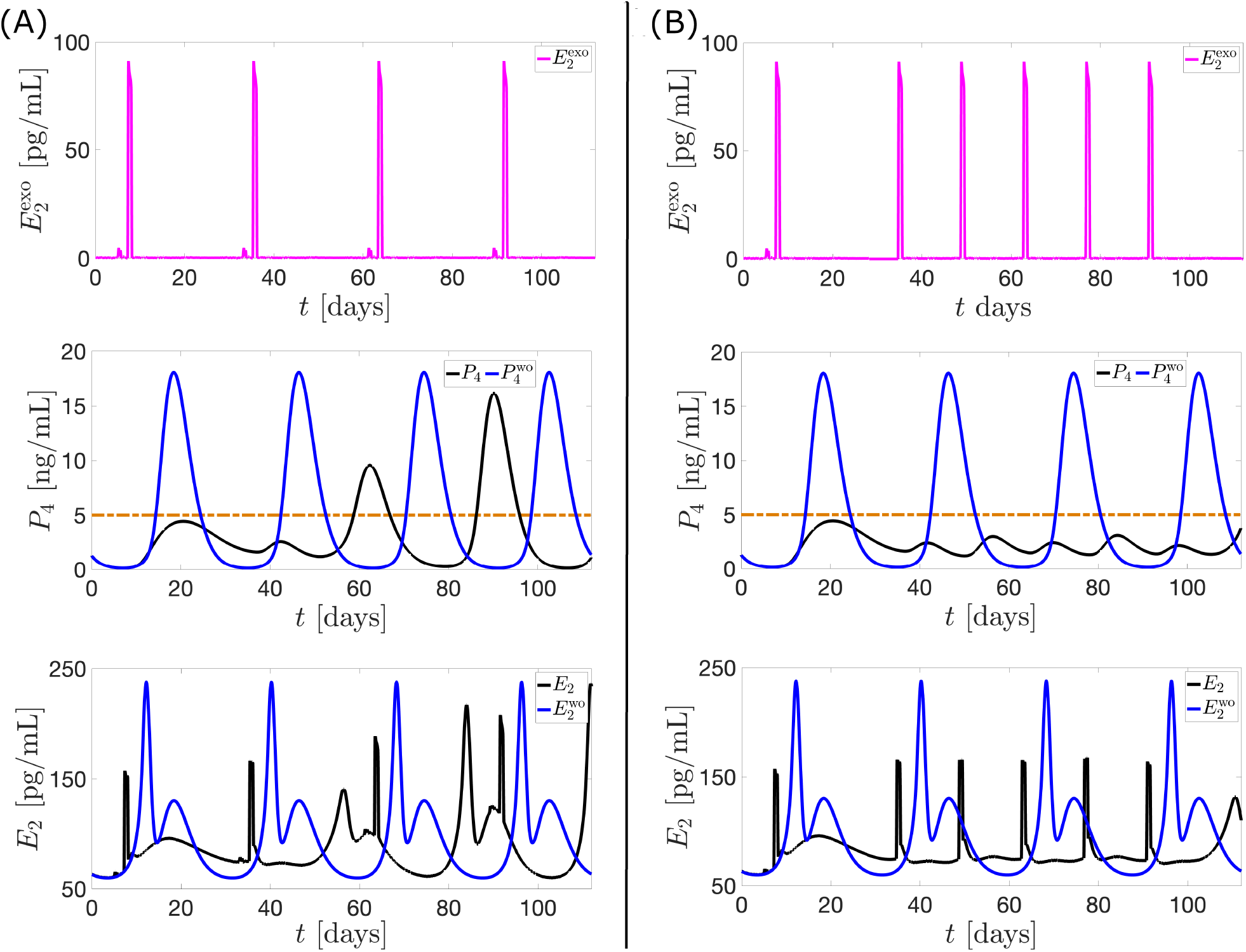
The black curve is the model output when multiple *u*_1_ (magenta curve) is applied. *H*^wo^ (blue curve) denotes the hormone model output without the influence of *u*_1_. In panel (A), the application of *u*_1_ from day 0 to day 112 in equal intervals is unable to sustain anovulation. Panel (B) shows a scheme for administration of multiple *u*_1_ which continuously blocks ovulation.

Hormonal contraceptive benefits go beyond contraception. Noncontraceptive advantages include: lower recurrence of endometriotic cyst; reduced risk of endometrial, ovarian, and colorectal cancer [46]; and reduction of anterior cruciate ligament (ACL) injury [47]. ACL injury is a prevalent injury in sports [48] and occurs at two to eight times higher rate in females than in males [49]. Aside from the great financial cost of treatment, 31% of patients had moderate to severe disability in walking, 44% had moderate to severe disability in daily functional activities, and 77% could not return to their pre-injury sports level [48, 50]. The paper [49] reported that ACL injuries in female athletes are significantly greater during the ovulatory phase and studies [47, 51, 52] suggested that oral contraceptive users have an almost 20% decreased risk for ACL injury. The current study may provide guidance on manipulating the menstrual cycle to prevent this type of injury. In particular, the optimal control approach may be used to investigate dosing regimen of contraceptives that eliminate substantial rise in estrogen levels during the cycle. This is because an increase in estrogen is presumed to decrease ACL’s tensile strength [53]. Estrogen also decreases motor skills, reducing the normal neuromuscular protective mechanisms of the knee [53].

## CONCLUSION

This study employs a menstrual cycle model which correctly predicts the pituitary and ovarian levels throughout a normal cycle and reflects the decrease in maximum hormone levels caused by exogenous estrogen and/or progesterone. Optimal control results show that a significant reduction in the dosage of exogenous estrogen and/or progesterone may induce anovulation. Furthermore, combination therapy lower doses even more. Simulations also show the effectivity of the administration of exogenous estrogen in the mid follicular phase.

Through the years, the reduction of exogenous estrogen and progesterone doses in contraceptives is done to decrease risks for adverse effects such as thrombosis and myocardial infarction. Hence, results presented in this paper may give clinicians guidance on conducting experiments about optimal treatment regimen causing anovulation. Because the model output cycle length of 28.05 days is a good approximation to the 28-day data cycle length, the model may be also be used to explore how treatments change the period of the menstrual cycle. Finally, this work may be refined by including absorption and clearance rates of the exogenous hormones to attain more realistic dosing schedules and allow investigation into effects of specific drugs. Early optimal control results in this study exhibit noticeable oscillations. By several adjustments to the objective function, we were able to smoothen the control. To further lessen fluctuations in the optimization outcome, usage of other numerical techniques like polyhedral active set algorithm or introduction of dynamic equation to the objective function may be explored. This work investigated optimization of dosing strategy over one cycle primarily due to the considerable amount of time it takes to obtain results. Therefore, optimal control for at least two cycles may be examined using more cost-effective numerical methods.

## ACKNOWLEDGEMENTS

We thank Dr. James F. Selgrade for the discussions which helped in analysis of results and improvement of the manuscript.

## APPENDIX

**TABLE V:** Model parameter symbols, descriptions, values, units, and references.

| Parameter | Description | Value | Unit | Reference |
| --- | --- | --- | --- | --- |
| $k_{LH}$ | $LH$ release to the blood rate | 0.9661 | $day^{-1}$ | estimated |
| $\alpha_{LH}$ | $LH$ clearance rate | 14.0 | $day^{-1}$ | [11] |
| $V_{0,LH}$ | $LH$ non-induced max synthesis rate | 550.03 | $IUday^{-1}$ | estimated |
| $V_{1,LH}$ | $LH$ max synthesis rate | 3329.19 | $IUday^{-1}$ | estimated |
| $km_{LH}$ | $E_2$ value at half-saturation | 136.05 | $pgmL^{-1}$ | estimated |
| Constant for inhibition of |  |  |  |  |
| $ki_{LH,P}$ | $LH$ synthesis | 6.78 | $ngmL^{-1}$ | estimated |
| $c_{LH,E}$ | $LH$ release to the blood | 0.0060 | $mLpg^{-1}$ | estimated |
| $c_{FSH,E}$ | $FSH$ release | 0.0151 | $\left(\frac{mL}{pg}\right)^2$ | estimated |
| $Ki_{FSH,Inh}$ | $FSH$ synthesis | 16.83 | $IUmL^{-1}$ | estimated |
| $w^*$ | $FSH$ synthesis | 9.21 | $IUmL^{-1}$ | estimated |
| $q^*$ | $RcF$ | 5.11 | $IUmL^{-1}$ | estimated |
| constant for stimulation of |  |  |  |  |
| $c_{LH,P}$ | $LH$ release to the blood | 1.98 | $mLng^{-1}$ | estimated |
| $c_{FSH,P}$ | $FSH$ release | 52.31 | $mLng^{-1}$ | estimated |
| $V_{FSH}$ | $FSH$ maximal synthesis rate | 294.90 | $IUday^{-1}$ | estimated |
| $\alpha_{FSH}$ | $FSH$ clearance rate | 8.21 | $day^{-1}$ | [11] |
| $k_{FSH}$ | $FSH$ release to the blood rate | 14.59 | $day^{-1}$ | estimated |
| $\tau$ | Time scale controlling $Inh$ concentration and $FSH$ synthesis | 1.5 | $days$ | [11] |
| $v$ | Blood volume | 2.50 | $L$ | [11] |
| Follicles transition factor from |  |  |  |  |
| $b$ | inactive to $RcF$ | 0.0453 | $\mu g(IUday)^{-1}$ | estimated |
| $c_2$ | $RcF$ to $GrF$ | 0.0577 | $(L/IU)^\alpha/day$ | estimated |
| $c_3$ | $GrF$ to $DomF$ | 0.0170 | $L/(IUday)$ | estimated |
| $c_4$ | $DomF$ to ovulation | 1.14 | $(L/IU)^\gamma/day$ | estimated |
| $d_1$ | $Sc_1$ to $Sc_2$ | 0.7537 | $day^{-1}$ | estimated |
| $d_2$ | $Sc_2$ to $Lut_1$ | 0.6866 | $day^{-1}$ | estimated |
| $k_1$ | $Lut_1$ to $Lut_2$ | 0.6699 | $day^{-1}$ | estimated |
| $k_2$ | $Lut_2$ to $Lut_3$ | 0.6388 | $day^{-1}$ | estimated |
| $k_3$ | $Lut_3$ to $Lut_4$ | 0.9191 | $day^{-1}$ | estimated |
| $c_1$ | $RcF$ stage follicles growth factor | 0.1036 | $LIU^{-1}day^{-1}$ | estimated |
| $k_4$ | $Lut_4$ stage degradation factor for corpus lutea | 1.88 | $day^{-1}$ | estimated |
| $\alpha$ | $LH$ exponent for promotion from $RcF$ to $GrF$ | 0.9505 | | estimated |
| $\gamma$ | $LH$ exponent for transition from $DomF$ to ovulation | 0.1615 | | estimated |
| $e_0$ | Min $E_2$ level | 57.60 | $pgmL^{-1}$ | estimated |
| Contribution factor of |  |  |  |  |
| $e_1$ | $GrF$ to $E_2$ | 0.0269 | $L^{-1}$ | estimated |
| $e_2$ | $DomF$ to $E_2$ | 0.4196 | $L^{-1}$ | estimated |
| $e_3$ | $Lut_4$ to $E_2$ | 0.4923 | $L^{-1}$ | estimated |
| $p_1$ | $Lut_3$ to $P_4$ | 0.0032 | $kL^{-1}$ | estimated |
| $p_2$ | $Lut_4$ to $P_4$ | 0.1188 | $kL^{-1}$ | estimated |
| $h_1$ | $DomF$ to $Inh$ | 0.0193 | $IUmL^{-1}\mu g^{-1}$ | estimated |
| $h_2$ | $Lut_3$ to $Inh$ | 0.0159 | $IUmL^{-1}\mu g^{-1}$ | estimated |
| $h_3$ | $Lut_4$ to $Inh$ | 0.0119 | $IUmL^{-1}\mu g^{-1}$ | estimated |
| $h_0$ | Min $Inh$ level | 0.6606 | $IUmL^{-1}$ | estimated |

\* added to the Margolskee model.

